# An insulin, AMPK, and steroid hormone-mediated metabolic switch regulates the transition between growth and diapause in *C. elegans*

**DOI:** 10.1101/323956

**Authors:** Sider Penkov, Bharath Kumar Raghuraman, Cihan Erkut, Jana Oertel, Roberta Galli, Eduardo Jacobo Miranda Ackerman, Daniela Vorkel, Jean-Marc Verbavatz, Edmund Koch, Karim Fahmy, Andrej Shevchenko, Teymuras V. Kurzchalia

## Abstract

The balance between growth and quiescence depends on the global metabolic state. The dauer larva of *C. elegans,* a developmentally arrested stage for survival under adverse environment, undergoes a major metabolic transition. Here, we show that this switch involves the concerted activity of several regulatory pathways. Whereas the steroid hormone receptor DAF-12 controls dauer morphogenesis, the insulin pathway maintains low energy expenditure through DAF-16/FoxO, which also requires AAK-2/AMPKα. DAF-12 and AAK-2 separately promote a shift in the molar ratios between competing enzymes at two key branch points within the central carbon metabolic pathway. This way, carbon atoms are diverted from the TCA cycle and directed to gluconeogenesis. When both AAK-2 and DAF-12 are suppressed, the TCA cycle is active and the developmental arrest is bypassed. Hence, the metabolic status of each developmental stage is defined by stoichiometric ratios within the constellation of metabolic enzymes and controls the transition between growth and quiescence.

## Introduction

Throughout their life cycle, organisms alternate between states of high and low metabolic activity. In some cases, not only the intensity but the whole mode of metabolism changes, for instance during the transition from growth to quiescence. Usually, growing organisms have highly active mitochondria and intensive oxidative phosphorylation (OXPHOS), whereas during metabolic quiescence they shift to glycolysis and associated gluconeogenesis [1, 2]. Most early embryos use glycolytic metabolism and only shift to OXPHOS during later phases of development [3]. This metabolic shift is also observed during the differentiation of neurons and stem cells [4, 5]. The opposite change, from OXPHOS to aerobic glycolysis, is seen in cancer cells exhibiting “Warburg” metabolism [6]. Despite their importance, however, we still have a limited understanding of mechanisms controlling these global metabolic transitions.

Entry of the nematode *Caenorhabditis elegans* into diapause is an excellent model in which to study these metabolic transitions. In response to harsh environmental conditions, *C. elegans* interrupts its reproductive life cycle, stops growing, and forms a specialized, developmentally arrested third larval stage called a dauer (enduring) larva [7]. The body of dauer larvae is morphologically adapted to harsh conditions. Its diameter is reduced, its body coated with a tight cuticle, and its pharynx sealed [7]. Most importantly, the metabolism of dauers differs substantially from that of the reproductive L3 larvae. Since they do not feed, they rely on stored energy reserves [7]. To restrict the depletion of these reserves, dauer larvae enter a hypometabolic mode via a dramatic rearrangement of anabolic and catabolic pathways [2,8–13]. In this “stand-by” mode, energy consumption, heat production, aerobic respiration and TCA cycle activity are significantly reduced. The production of cofactors required for anabolic reactions such as NADPH is also minimized [14]. In addition, the glyoxylate shunt and gluconeogenesis are used to generate carbohydrates from reserve lipids [2,8–13]. In this state, dauers can survive for months without nutrition.

The process of dauer formation is controlled by Daf genes (from <u>da</u>uer <u>f</u>ormation). Whereas Daf-<u>c</u> mutants <u>c</u>onstitutively undergo dauer arrest, Daf-<u>d</u> mutants are <u>d</u>efective in forming dauer larvae. Genetic analysis of Daf mutants has revealed that dauer formation is governed by guanylyl cyclase, TGF-β-like, insulin-like and steroid hormone signaling pathways [7, 15] (**Fig. 1A**). In response to changes in population density (sensed through dauer-inducing pheromones) and altered energetic metabolism (signaled by insulin-like peptides), the guanylyl cyclase, TGF-β and insulin-like pathways converge on two transcription factors: the FoxO member DAF-16 and the nuclear hormone receptor DAF-12, both encoded by Daf-d genes that are essential for dauer formation. DAF-16 is negatively regulated by the insulin receptor homolog DAF-2 in response to stimulation by insulin-like peptides [16–18]. DAF-12, on the other hand, is regulated by steroid hormones, called dafachronic acids (DAs), synthesized by the cytochrome P450 enzyme DAF-9 when the population density is low [19–22]. DAs bind to DAF-12 and suppress its dauer-promoting activity [19,23,24]. DAF-16 and DAF-12 stimulate each other but also have their own downstream programs (**Fig. 1A**) [25, 26]. The interplay between these factors determines whether worms enter diapause: when both are activated, dauer formation is induced. In addition, a germline-mediated crosstalk between DAF-16 and DAF-12 is essential for adult longevity [27]. Although many transcriptional, as well as metabolic targets, of DAF-16 and DAF-12 have been elucidated in the context of diapause and longevity [14,28–37], fundamental questions remain about how these transcription factors interact to control the metabolism and what is the impact of the metabolic switch on the growth and development.

**Figure 1.**
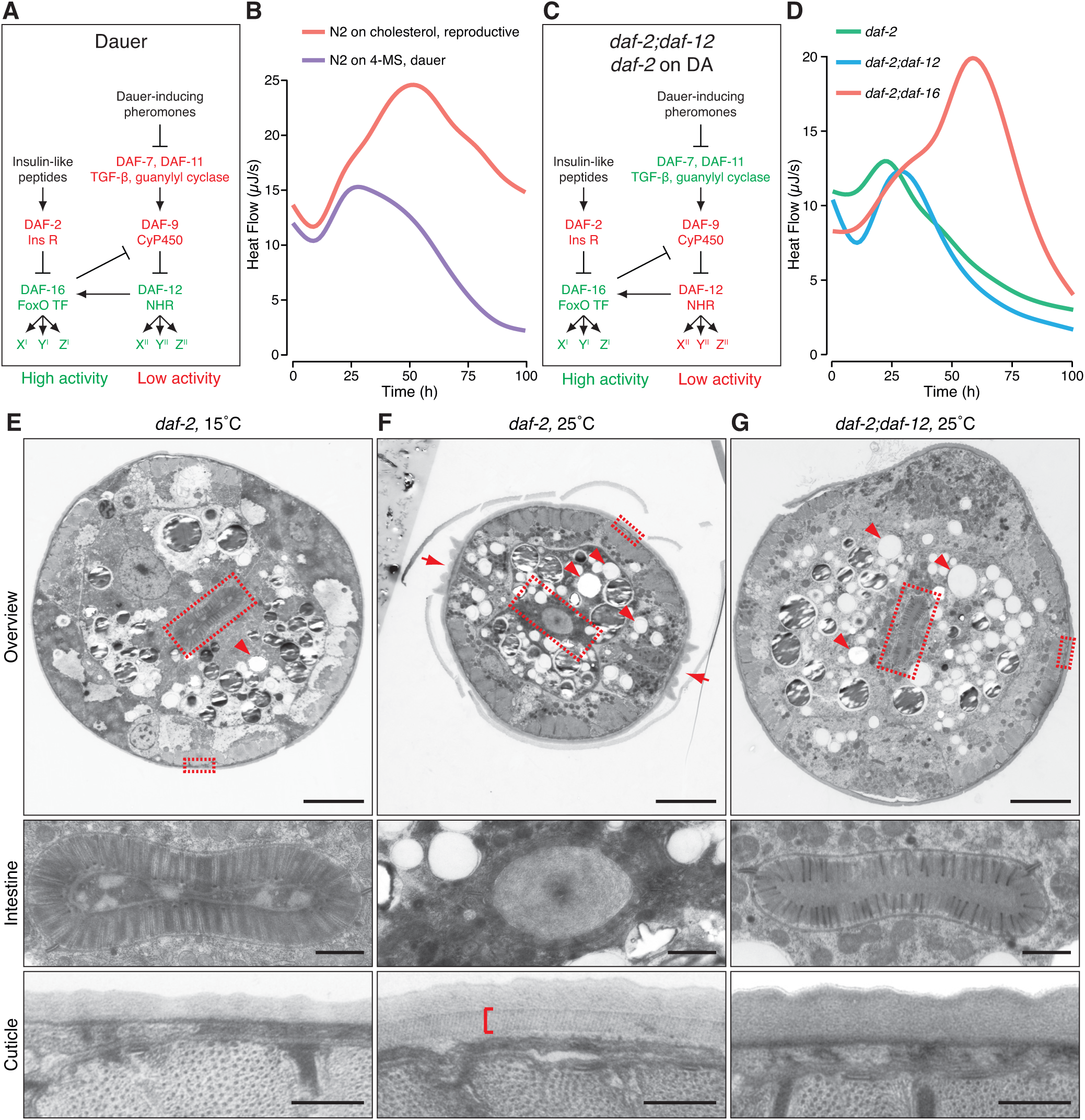
DAF-16 mediates the switch to low metabolic rate in dauer formation but does not directly induce dauer morphogenesis. **(A)** Signaling pathways in dauer formation. In the absence of insulin-like peptides, DAF-2 is suppressed, leading to activation of DAF-16. Dauer-inducing pheromones inhibit the TGF-β and guanylyl cyclase pathways, and this lowers the production of DA by DAF-9, resulting in activated DAF-12. DAF-16 stimulates DAF-12 by inhibiting DAF-9. DAF-12 is also capable of activating DAF-16. DAF-16 and DAF-12 control different subsets of genes designated X′Y′Z′ and X¢¢Y¢¢Z¢¢, respectively. **(B)** Continuous measurement of the heat dissipated in unit time (heat flow) by wild-type (N2) worms undergoing reproductive growth on cholesterol or dauer formation on 4-MS. **(C)** Formation of L3 arrested larvae of *daf-2;daf-12* or DA-fed *daf-2* at restrictive temperature. The reduction of DAF-2 activity leads to activation of DAF-16. DAF-16 inhibits DAF-9 activity but this effect is neutralized by the *daf-12* mutation or the addition of exogenous DA. Thus, DAF-16 is active but DAF-12 is not. **(D)** Heat flow of *daf-2, daf-2;daf-12* and *daf-2;daf-16* grown at 25°C. **(E)** Transmission electron micrograph of a cross-section of a growing L3 larva of *daf-2* grown at 15°C. Upper panel: an overview of the body organization. The gut lumen is elongated and lined by multiple microvilli (central panel, large rectangle in the upper panel). The structure of the cuticle is displayed on the lower panel, corresponding to the small rectangle in the upper panel. Note the absence of a striated circular layer. **(F)** Electron micrograph of a *daf-2* dauer larva grown at 25°C. The upper panel shows radial constriction of the body, extensive accumulation of lipid droplets (arrowheads) and dauer-specific alae (arrows). The gut lumen is rounded, microvilli are almost absent (central panel, large rectangle in upper panel). The cuticle possesses a striated layer (lower panel and indicated by a bracket, small rectangle in the upper panel). **(G)** Electron micrograph of a *daf-2;daf-12* arrested larva grown at 25°C. The body is not radially constricted but displays extensive accumulation of lipid droplets (upper panel, arrowheads). Alae are absent (upper panel). The gut lumen (central panel, large rectangle in upper panel) and the cuticle (lower panel, small rectangle in upper panel) resemble the ones displayed by L3 larvae in (Fig. 1E). 4-MS – 4-methylated sterol; DA – dafachronic acid. Panels **B** and **D** show representative diagrams from 4 experiments with 1-4 technical replicates. Scale bars in **E**, **F** and **G** correspond to 5 *μ*m (upper panels), 1 *μ*m (central panels) and 0.5 *μ*m (lower panels); representative images of at least five animals per condition.

Here, we show that during dauer formation, the metabolic mode, consisting of two separately regulated modules, dictates the state of development. The insulin pathway has a dual effect that requires AMP-activated protein kinase (AMPK) activity. On one hand, it maintains low catabolism (first module). On the other, it acts together with the steroid hormone pathway to inhibit the TCA cycle and promote gluconeogenesis (second module). Simultaneous inactivation of the AMPK and steroid hormone pathways leads to a switch from gluconeogenesis to an active TCA cycle via tight control of the molar ratios of competing enzymes. This metabolic transition is a prerequisite for the the organism to enter into reproductive growth and is conserved in long lived adults with reduced insulin signaling. Moreover, the state of metabolism that is dictated by the stoichiometric proportions between enzymes of the central carbon metabolism can be used to predict the transition from growth to quiescence.

## Results

### DAF-16 induces a switch to low metabolic rate whereas DAF-12 controls dauer morphogenesis

To distinguish which signaling pathways control metabolism in the dauer state, we investigated the metabolic activities of wild-type dauers, as well as of mutants of key dauer regulatory factors. Owing to an overall lower metabolic rate, dauer larvae have substantially diminished heat production compared to other larval stages [10]. For that reason, we first compared the heat flow produced by wild-type worms undergoing reproductive development or dauer formation from the L1 larval stage onwards using time-resolved isothermal microcalorimetry. To induce a synchronous dauer formation, we grew worms on 4-methylated sterol (4-MS) which blocks the production of DAs [26]. After an initial increase of heat flow by both groups, the two trends diverged after about 24 hours, increasing further in worms in the reproductive mode, while decreasing in animals that underwent dauer formation (**Fig. 1B**). A similar trend was observed in the TGF-β Daf-c mutant *daf-7*, which forms dauer larvae but reproduces when DA is added [19, 38] (**Supplementary Fig. S1A**).

Next, we set out to determine how the insulin and steroid hormone pathways contribute to the switch. To disentangle the pathways, we chose conditions under which DAF-16 is active but DAF-12 not. We made use of a group of Daf-c alleles of *daf-2*, designated class II, that are not fully suppressed by Daf-d mutations of *daf-12* or by addition of DAs. One such allele is *daf-2(e1370)*. Worms bearing this mutation reproduce at the permissive temperature of 15°C, whereas at the restrictive temperature of 25°C, they form dauers. We compared this strain to a double mutant *daf-2(e1370);daf-12(rh61rh411)* or DA-treated *daf-2(e1370)* that arrest the development at an L3-like larval stage at 25°C due to DAF-16 activity (**Fig. 1C**) [7,19,25]. The metabolism and morphology of these larvae have not been characterized in detail. Unlike *daf-7* on DA, at 25°C *daf-2;daf-12* and *daf-2* on DA shifted the metabolic mode to low heat production after 24 hours (**Fig. 1D** and **Supplementary Fig. S1A**). In contrast, a double mutant *daf-2(e1370);daf-16(mu86)* that undergoes reproductive development at 25°C [25] displayed high heat production (**Fig. 1D**). Thus, activation of DAF-16, but not of DAF-12, mediates the switch to low metabolic activity during dauer formation.

We further asked whether DAF-16 or DAF-12 determines the morphology of dauer larvae and whether metabolic state and morphology are interconnected. Our previous studies indicated that DAF-12 can induce morphological features of dauer larvae in the absence of DAF-16 [26]. However, it was not clear whether activation of DAF-16 alone could promote dauer morphology. To test this, we performed electron microscopy on *daf-2;daf-12* and DA-treated *daf-2* larvae grown at 25°C. Interestingly, similar to L3 larvae, they had a large body diameter, an elongated gut lumen with long, densely packed microvilli, and lacked characteristic dauer features such as alae and a striated layer (**Fig. 1E, F, G** and **Supplementary Fig. S1B**) [39]. However, similar to dauers, *daf-2;daf-12* and DA-fed *daf-2* animals deposited numerous lipid droplets (LDs) **(Fig. 1G** and **Supplementary Fig. S1B**). Thus, DAF-12 controls dauer morphogenesis, whereas DAF-16 has no direct influence on this process but appears to affect the dauer-associated metabolic changes that culminate in low metabolic rate and high LD accumulation.

### DAF-16 controls catabolism and, together with DAF-12, promotes a shift from TCA cycle-driven metabolism to gluconeogenesis

Cells produce heat almost exclusively through catabolic reactions [40]. Thus, low heat production in dauers indicated that they have decreased catabolism. To determine which pathways regulate this process, we compared the amounts of heat that *daf-2* dauers and *daf-2;daf-12* L3-like larvae produce after entering this arrested state. Food was omitted to exclude heat generation by bacteria. As seen in **Figure 2A**, *daf-2* dauers and *daf-2;daf-12* larvae generated similar amounts of heat, suggesting that DAF-16 does not require DAF-12 activity to regulate the energy expenditure. We next asked how loss of DAF-16 activity would influence the metabolic rate. For inactivation of DAF-16, we used *daf-16(mu86)* mutants grown on 4-MS. Under these conditions, DAF-12 promotes the dauer program, but DAF-16 is absent and worms arrest as dauer-like animals [26] (**Fig. 2B**). Compared to wild-type dauers, 4-MS treated *daf-16* animals displayed higher heat production in their arrested state (**Fig. 2C**), suggesting that DAF-16 suppresses metabolic rate and the catabolism of energy stores of dauers.

**Figure 2.**
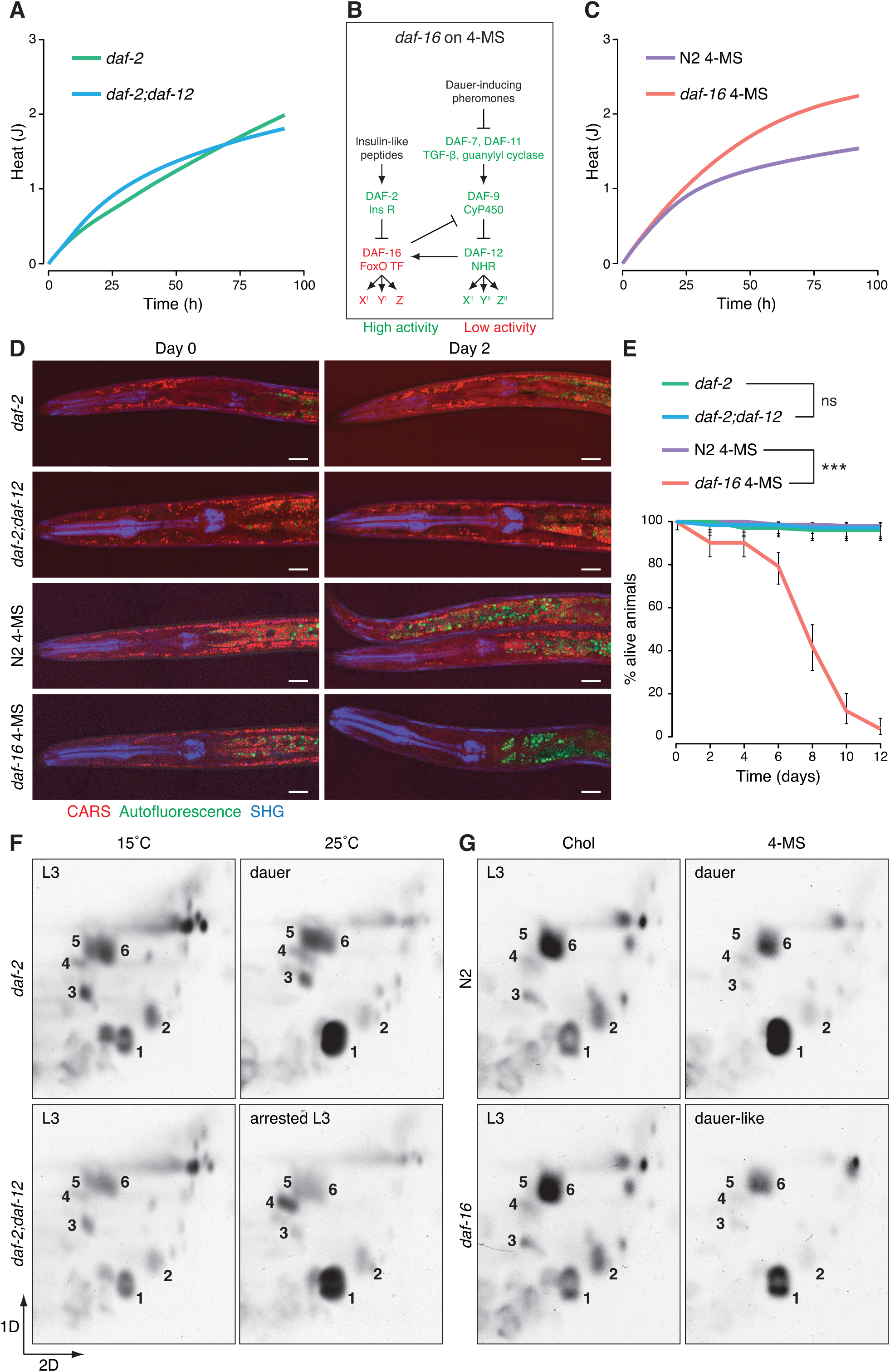
DAF-16 alone determines the energy expenditure and lifespan of dauer larvae and, together with DAF-12, the switch to gluconeogenesis. (**A**) Continuous measurement of the cumulative heat dissipation by *daf-2* dauers and *daf-2;daf-12* arrested L3 larvae grown at 25°C in the period after the developmental arrest is completed. (**B**) Formation of dauer-like larvae of *daf-16* on 4-MS. DAF-12 is activated due to a lack of substrates for DA synthesis; however, DAF-16 activity is absent due to the mutation in the *daf-16* locus. (**C**) Cumulative heat dissipation by wild-type (N2) dauers and *daf-16* dauer-like larvae grown on 4-MS in the period after the developmental arrest is completed. (**D**) CARS microscopy of lipid droplets (red) in *daf-2* dauers and *daf-2;daf-12* arrested L3 larvae grown at 25°C, and wild-type (N2) dauers and *daf-16* dauer-like larvae grown on 4-MS in the period after the developmental arrest is completed. Note that the lysosome-related organelles (green autofluorescence) are not a source of CARS signal under the conditions used. (**E**) Survival rate of *daf-2* dauers and *daf-2;daf-12* arrested L3 larvae grown at 25°C, and wild-type (N2) dauers and *daf-16* dauer-like larvae grown on 4-MS in buffer. (**F**) 2D-TLC of ^14^C-acetate-labelled metabolites from *daf-2* dauers and *daf-2;daf-12* arrested larvae grown at 25°C compared to growing L3 larvae of the same strains grown at 15°C. Note the accumulation of trehalose (1) in the arrested stages, indicating a switch from TCA cycle to gluconeogenesis. (**G**) 2D-TLC of ^14^C-acetate-labelled metabolites from wild-type (N2) dauers and *daf-16* dauer-like larvae grown on 4-MS compared to growing L3 larvae of the same strains grown on cholesterol. 4-MS – 4-methylated sterol, CARS – Coherent Anti-Stokes Raman Scattering, SHG – Second Harmonic Generation. In (**A**) and (**C**), representative diagrams of at least 2 experiments with 3 replicates. In (**D**), representative images of at least 6 animals, scale bars – 10 *μ*m. In (**E**), data is represented as means ± 95% confidence intervals of 3 experiments with 3 replicates. *** significance of p<0.001; ns - no significant difference determined by log-rank test. In (**F** and **G**), representative images from at least 2 experiments. 1 – trehalose, 2 – glucose, 3 – glutamate, 4 – glycine/serine, 5 – glutamine, 6 – alanine/threonine.

We therefore monitored the breakdown of lipids, sugars and amino acids in *daf-2;daf-12* and *daf-16* on 4-MS. Because food was omitted, only the catabolism of internal energy reserves would account for any observed change. Storage triglycerides (TGs) were visualized by Coherent Anti-Stokes Raman Scattering (CARS) microscopy of LDs and by thin-layer chromatography (TLC). *daf-2* dauers and *daf-2;daf-12* arrested larvae, as well as wild-type dauers on 4-MS, efficiently conserved their TGs over time (**Fig. 2D and Supplementary Fig. S2A**). In contrast, *daf-16* larvae on 4-MS were depleted of TGs only after two days (**Fig. 2D and Supplementary Fig. S2A**). Phospholipids were preserved in all animals, suggesting that no major degradation of membranes occurred (**Supplementary Fig. S2A**). Furthermore, sugars and amino acids were maintained at high levels in wild-type dauers on 4-MS, *daf-2*, and *daf-2;daf-12* larvae, but were rapidly degraded in *daf-16* on 4-MS (**Supplementary Fig. S2A** and **S2B**). These results show that, in the absence of DAF-16, catabolism becomes misregulated and the energy depot is dramatically depleted.

The above findings suggest that faster depletion of energy reserves might reduce survival in *daf-16* larvae. Indeed, the viability of these animals declined rapidly and they perished after 12 days, while almost 100% of dauers and *daf-2;daf-12* arrested larvae remained viable (**Fig. 2E**). Thus, DAF-16 regulates the survival of dauer larvae by controlling energy expenditure.

The switch to a lower catabolic rate in dauers is accompanied by a shift from TCA cycle-driven metabolism to gluconeogenesis, leading to accumulation of the disaccharide trehalose [9]. To determine whether DAF-16 or DAF-12 is responsible for this transition, we used 2D-TLC to trace the metabolism of ^14^C-radiolabeled acetate. Carbon atoms of acetate are only incorporated into trehalose if the glyoxylate shunt and the gluconeogenesis are active [9]. This labeling strategy mimics the usage of endogenous lipids as a carbon source for gluconeogenesis because both the lipid catabolism and the external acetate provide acetyl-CoA that enters the TCA or the glyoxylate cycle. As shown before [9], *daf-2* dauers at 25°C displayed stronger accumulation of labelled trehalose than L3 larvae at 15°C (**Fig. 2F**). High incorporation of acetate into trehalose was also observed in *daf-2;daf-12* arrested L3 larvae (**Fig. 2F**). Thus, activation of DAF-16 is sufficient to trigger gluconeogenesis. Surprisingly, we also detected higher levels of labelled trehalose in *daf-16* mutants cultured on 4-MS, suggesting that DAF-12 can promote a gluconeogenic mode in the absence of DAF-16 (**Fig. 2G**). Together, our results demonstrate that DAF-16 alone maintains low catabolism, whereas DAF-16 and DAF-12 separately promote a shift from TCA cycle-driven metabolism to gluconeogenesis. In addition, the low catabolism and the gluconeogenic mode are independent metabolic modules that can be uncoupled under conditions of low DAF-16 but high DAF-12 activity.

### AAK-2 is required for the DAF-16-mediated metabolic switch, developmental arrest, and adult longevity

We postulated that under conditions of high DAF-16 but low DAF-12 activity (**Fig. 1C**), the disruption of a hypothetical factor required for the DAF-16-mediated metabolic switch could promote higher catabolism and prevent the gluconeogenic mode. Moreover, if the metabolic and developmental transition are coupled, one prediction would be that such an intervention may rescue the developmental arrest caused by DAF-16. Thus, it was of high importance to identify such a factor. The uncontrolled catabolism and mortality in *daf-16* on 4-MS were very similar to that observed in dauers with loss of activity of the AMPK α-subunit AAK-2 [41]. Thus, DAF-16 and AAK-2 may jointly control the metabolic state of dauers, making AMPK a potential candidate for this factor. We first asked whether *aak-2* mutant dauers lose TGs, sugars and amino acids similar to *daf-16* on 4-MS. We generated a *daf-2(e1370);aak-2(gt33)* strain harboring a large deletion in *aak-2*. At 25°C, almost all animals formed dauer larvae with typical dauer morphology (**Fig. 3A**). Curiously, although a previous study that used a strain *daf-2(e1370);aak-2(ok524)*, bearing a different deletion in *aak-2*, showed that at 25°C the animals spontaneously exited from dauer state and produced adults within five days [42], dauers of *daf-2(e1370);aak-2(gt33)* grown on a solid medium with ample food for five days at 25°C did not undergo spontaneous exit from dauer state. Almost all worms survived treatment with the detergent SDS, which is a hallmark of dauer larvae [43] (**Supplementary Fig. S3A** and **S3B**). The dauer-specific alae and striated layer of the cuticle were also preserved over time (**Fig. 3A**). Hence, using *daf-2(e1370);aak-2(gt33)* is a very suitable model to study the metabolic control in dauer state.

**Figure 3.**
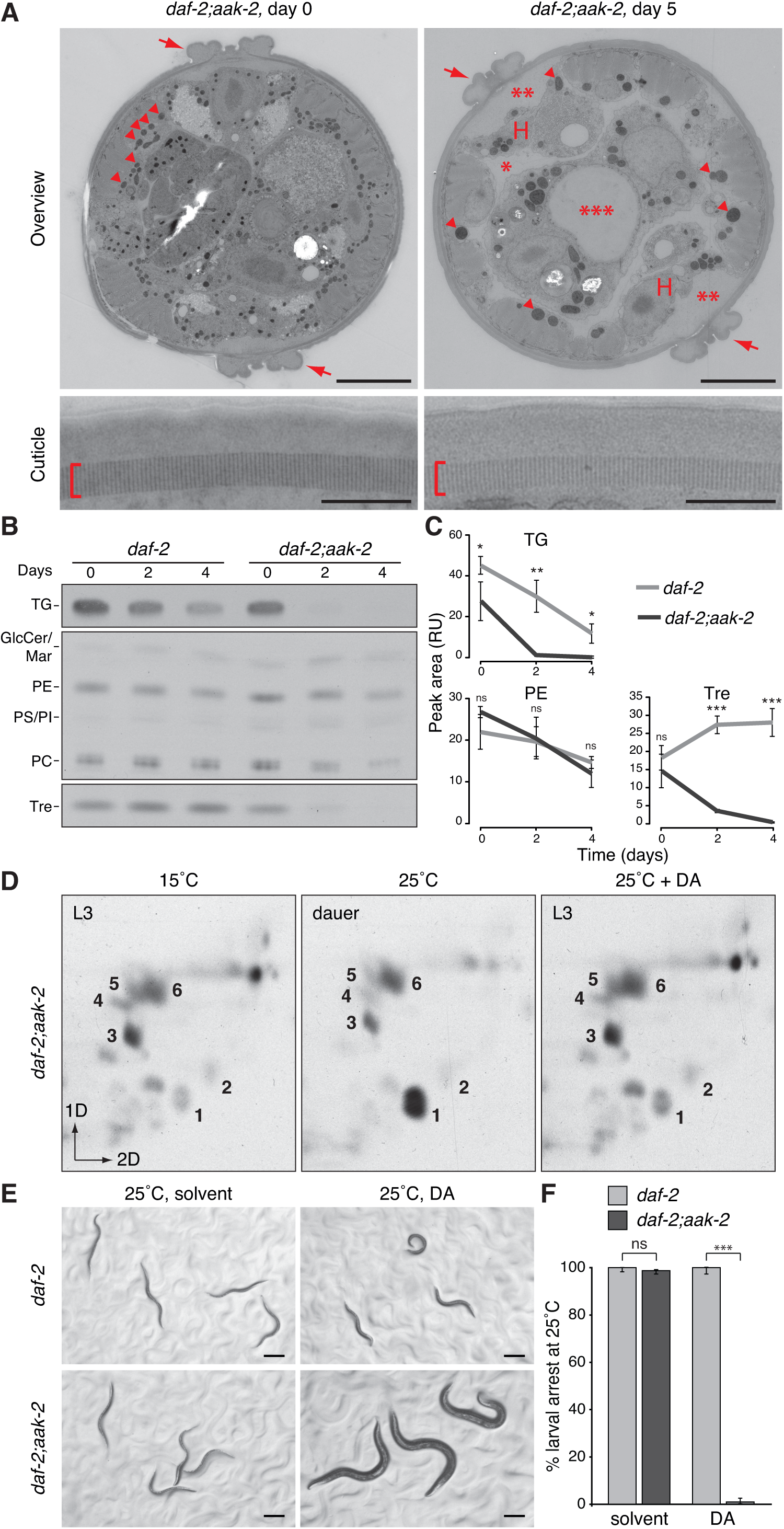
AAK-2 regulates the catabolism in dauer state and, in parallel to DAF-12, the gluconeogenic mode and developmental arrest. **(A)** Electron microscopy of *daf-2(e1370);aak-2(gt33)* collected on the day of dauer formation or after five days of incubation at 25 °C on ample food source. Alae (upper panels, arrows) and striated layer of the cuticle (lower panels, brackets) are present. Five day-old dauers show features of starvation: shrinkage of the hypodermis (H), expansion of the pseudocoelomic cavity (1 asterisk), formation of a cavity between the cuticle and the hypodermis (2 asterisks) and widening of the gut lumen (3 asterisks). In addition, the animals show much fewer mitochondria which are substantially enlarged (arrowheads). **(B)** TLC of ^14^C-acetate labelled TGs, phospholipids, and trehalose from *daf-2* and *daf-2;aak-2* dauers measured at different time points after the arrest. **(C)** Quantification of the band intensities in some of the compounds in (**B**) represented by peak area of the optical density. PE is used as a representative phospholipid. **(D)** 2D-TLC of ^14^C-acetate-labelled metabolites from *daf-2;aak-2* L3 larvae grown at 15°C, dauers grown at 25°C and L3 larvae obtained at 25°C in the presence of DA. **(E)** Micrographs of *daf-2* and *daf-2;aak-2* animals grown at 25°C in the presence or absence of DA. Inhibition of DAF-12 promotes reproductive growth in *daf-2;aak-2* but not in *daf-2*. **(F)** Quantification of the larval arrest in (**E**). In (**A**), Scale bars 5 *μ*m (upper panels) and 0.5 *μ*m (lower panels); representative images of at least 4 animals. In (**B** and **C**), TG – triglycerides, GlcCer – glucosylceramides, Mar – maradolipids, PE – phosphatidylethanolamine, PS – phosphatidylserine, PI – phosphatidylinositol, PC – phosphatidylcholine, Tre – trehalose, RU – relative units. In (**C**) data is represented as means ± SD of 2 experiments with 3 replicates. *** p<0.001; ** p<0.01; * p<0.1; ns - no significant difference determined by Student t-test. In (**D**), DA – dafachronic acid, 1 – trehalose, 2 – glucose, 3 – glutamate, 4 – glycine/serine, 5 – glutamine, 6 – alanine/threonine; representative images from at least 2 experiments. In (**E** and **F**) data is represented as means ± 95% confidence intervals of 3 experiments with 3 replicates. *** p<0.001; ns - no significant difference determined by one-way analysis of variance.

Electron micrographs suggested that after five days *daf-2;aak-2* dauers enter a state of starvation characterized by an extreme decrease of the cellular volume of the hypodermis, expansion of the body cavities and deterioration of mitochondria (**Fig. 3A**). In line with these observations, TGs and trehalose (**Fig. 3B** and **C**), as well as amino acids (**Supplementary Fig. S3C** and **S3D**) were rapidly depleted. Phospholipids were less affected (**Fig. 3B and C**). To test the cellular response to starvation in AMPK mutants, we monitored FIB-1, a small nucleolar ribonucleoprotein (snoRNP) whose localization to the nucleolus is reduced during starvation or loss of TOR activity [44]. Wild-type dauers isolated from overcrowded plates had nucleolar FIB-1 consistent with a non-starved state (**Supplementary Fig. S3E**). *daf-2* and *daf-2;aak-2* dauers, as well as *daf-2* DA-fed larvae, also displayed nucleolar FIB-1 shortly after the arrest (**Supplementary Fig. S3F, S3G and S3H**). This localization remained unchanged over time in *daf-2* dauers and *daf-2* DA-fed arrested L3 larvae (**Supplementary Fig. S3F and S3G**). In *daf-2;aak-2* dauers, however, FIB-1 formed granular structures in the nucleoplasm after four days and was almost completely dispersed in the nucleoplasm of cells after seven days (**Supplementary Fig. S3H**). Thus, *daf-16* and *aak-2* mutants have highly related phenotypes in terms of catabolism in dauer state and may act in the same pathway. Moreover, DAF-16 could prevent a TOR-dependent starvation response in the absence of DAF-12, but not of AAK-2, suggesting that AMPK is required for the DAF-16-mediated maintenance of the energy reserves.

Since the disruption of *aak-2* enhanced catabolism in *daf-2* dauers, we asked if it could also abolish the gluconeogenic mode. ^14^C-acetate labelling in *daf-2;aak*-2 at 25°C showed a pronounced gluconeogenic mode (**Fig. 3D**). Because DAF-12 could activate gluconeogenesis in the absence of DAF-16, we asked whether it could perform this activity also in the absence of AAK-2. Remarkably, when we inhibited DAF-12 in *daf-2;aak-2* by adding DA, the gluconeogenesis was abolished (**Fig. 3D**). Hence, AAK-2 fulfils the criteria for a factor required for the switch to low energy expenditure and to gluconeogenesis induced by DAF-16 when DAF-12 is inhibited. To determine whether AAK-2 is also necessary for the DAF-16 induced growth arrest, we monitored the development of *daf-2;aak-2* worms at 25°C with DA. Astoundingly, these worms completely bypassed dauer arrest and developed into adults (**Fig. 3E** and **F**). Thus, in *daf-2* mutants, AAK-2 is essential for the DAF-16 mediated growth and metabolic transition in the absence of DAF-12 activity.

An interaction between *daf-2* and *aak-2* has also been observed in the context of adult longevity: the lifespan extension characteristic for *daf-2* animals is fully suppressed by *aak-2* mutations [45]. Thus, the metabolic mode associated with increased longevity in adult *daf-2* mutants [46] could depend on AAK-2. To assess the gluconeogenesis, we labeled *daf-2* and *daf-2;aak-2* with ^14^C-acetate and grew them at 15°C until L4 stage to bypass dauer formation. From this point on, we either kept them at 15°C to maintain the DAF-2 activity high or shifted them to 25°C to suppress DAF-2. This temperature shift doubles the lifespan of *daf-2* adults compared to the wild-type worms [47]. 24 hours later, we extracted the metabolites and observed much higher accumulation of labelled trehalose in *daf-2* worms at 25°C as compared to 15°C (**Supplementary Fig. S3I**). *daf-2;aak-2* displayed lower incorporation of ^14^C-acetate into trehalose at both 15°C and 25°C in comparison to *daf-2.* This observation shows that AAK-2 is required for the full extent of the metabolic switch in *daf-2* adults. A small elevation of labelled trehalose in *daf-2;aak-2* at 25°C compared to 15°C (**Supplementary Fig. S3I**) suggests that in adults, DAF-16 could also promote gluconeogenesis to a very limited degree in an AAK-2-independent manner. Thus, the metabolic switch does not only determine dauer diapause, but also the lifespan of adults with reduced insulin signaling.

### Gluconeogenesis is turned on by a shift in the molar ratios of key metabolic enzymes

To gain insight into the molecular mechanism underlying the switch to gluconeogenesis and how AAK-2 and DAF-12 control it, we employed the LC-MS/MS method of MS Western [48] to quantify the absolute (molar) amount of 43 individual enzymes or subunits of enzymatic complexes involved in TCA cycle, glyoxylate shunt, glycolysis, gluconeogenesis and mitochondrial pyruvate metabolism. Molar abundance of each protein was determined by comparing individual abundances of several (typically, 2 to 5) quantitypic peptides with ^13^C, ^15^N-isotopically labeled peptide standards (**Supplementary Fig. S4A-F**). Standard peptides were concatenated into a protein chimera (**Supplementary Fig. S4G**) that was in-gel co-digested with target proteins separated by one-dimensional SDS PAGE from a whole animal lysate. The molar amount of chimera protein was referenced to the standard of BSA and quantified in the same LC-MS/MS experiment. MS Western quantification was highly concordant. Median coefficient of variation of molar abundances of proteins determined using alternative standard peptides was less than 10% (**Supplementary Fig. S4H**) with better than 0.99 Pearson coefficient of correlation between technical replicas (**Supplementary Fig. S4I**). The molar abundances of individual proteins were normalized to the total protein content in each animal lysate and could be directly compared between all biological conditions without metabolic or chemical labeling of target proteins.

We first analysed the enzyme levels in dauers (*daf-2* at 25°C) and L3 larvae (*daf-2* at 15°C). The 43 enzymes were detected in a wide range of 1 to nearly 160 fmol per μg of the total protein (**Fig. 4A, Supplementary Fig. S5** and **S6A**). Although a global metabolic perturbation would be expected, we found that the balance of molar abundances of members of different pathways between the two groups was not perturbed. However, in dauers the enzymes of glycolysis/gluconeogenesis were slightly more prevalent in respect to other pathways indicating enhanced gluconeogenesis (**Supplementary Fig. S6B**). Interestingly, in total, dauers were 1.7-fold more enriched in metabolic enzymes compared to L3 larvae, despite that dauer is a metabolically reduced stage (**Supplementary Fig. S6C**). Glycolysis/gluconeogenesis enzymes were enriched at the higher rate. (**Supplementary Fig. S6D**). Hence, the overall architecture of metabolic network was preserved in both developmental conditions, while the metabolic switch was executed by fine-tuning its directionality.

**Figure 4.**
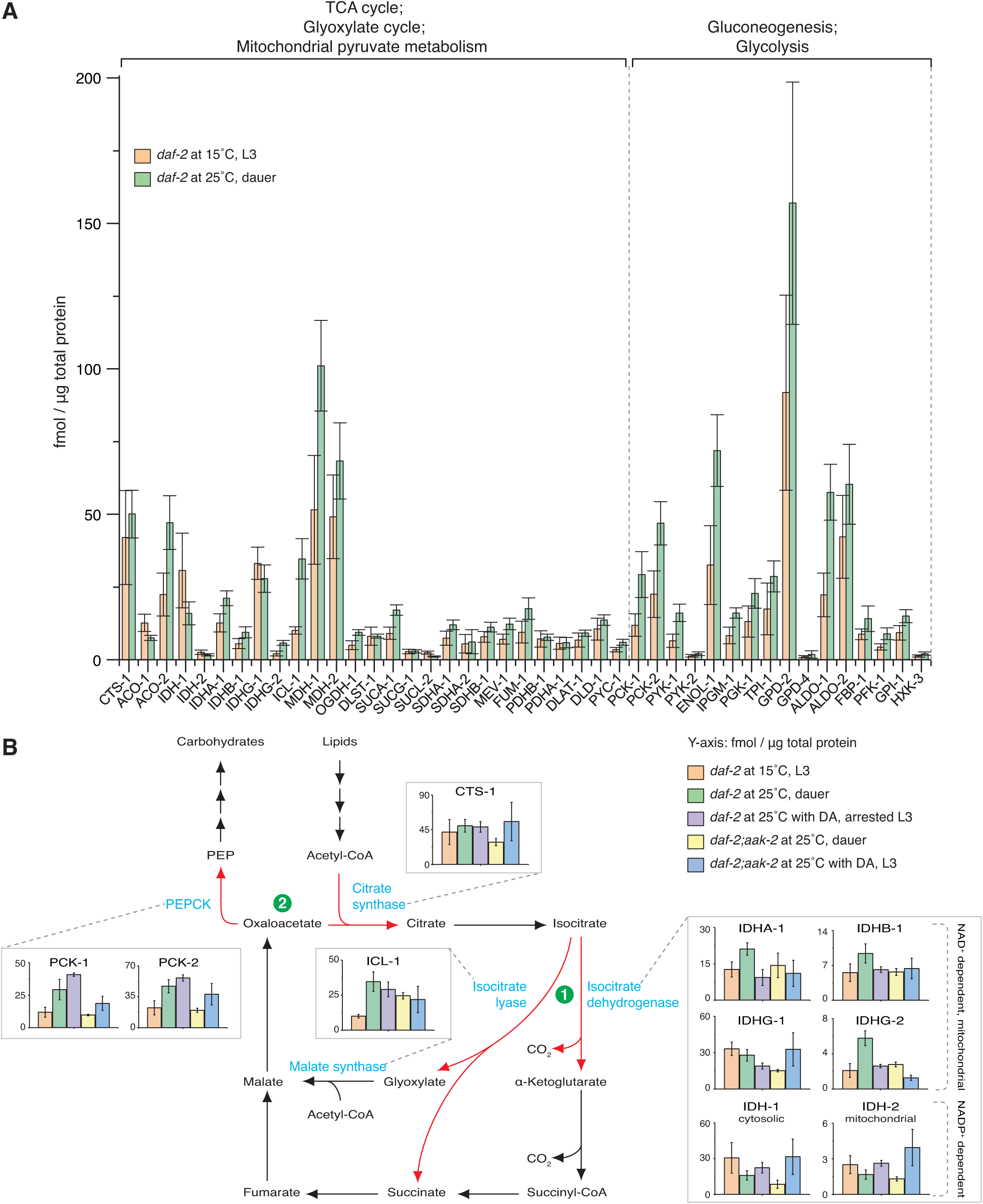
The metabolic switch is achieved through regulation of enzymes that work on branching points between competing pathways. (A) Absolute quantification of 43 enzymes of the TCA and glyoxylate cycle, mitochondrial pyruvate metabolism, gluconeogenesis, and glycolysis in *daf-2* dauers grown at 25°C compared to *daf-2* L3 larvae at 15°C. (**B**) Schematic representation of the pathway that converts lipids to carbohydrates with absolute quantification of the enzymes that operate at the branching points. Red arrows and green circles represent the competing reactions at the point of divergence between (1) TCA and glyoxylate pathway and (2) the recycling of oxaloacetate into the TCA/glyoxylate cycle or its entry into gluconeogenesis. In all panels, data is represented as means ± standard deviation (S.D.) of 3 biological replicates with 2 technical replicates each.

To understand metabolic switch mechanism, we reconstructed the pathway that converts lipid-derived acetyl-CoA to carbohydrates via glyoxylate shunt and gluconeogenesis (**Fig. 4B, Supplementary Fig. S5** and **S6A**). We focused on several reactions serving as branching points or “metabolic turnouts”. The first set of reactions determines whether acetyl-CoA, via isocitrate, will enter the TCA or the glyoxylate cycle (**Fig. 4B** and **Supplementary Fig. S5)**. We observed 3.5-fold upregulation of the glyoxylate cycle enzyme ICL-1 in dauers compared to L3 larvae (**Fig. 4B** and **Supplementary Fig. S5**). This was consistent with previous studies [29, 46] and suggested that dauers have higher glyoxylate pathway activity. Thus, they convert isocitrate to malate and succinate without losing carbon atoms in the TCA cycle via decarboxylation (**Fig. 4B** and **Supplementary Fig. S5**). Whether these carbon atoms are used for gluconeogenesis depends on the reactions at the second branching point. They determine whether the oxaloacetate produced downstream of the glyoxylate pathway is recycled by the citrate synthase or converted to phosphoenolpyruvate (PEP) by phosphoenolpyruvate carboxykinase (PEPCK) (**Fig. 4B** and **Supplementary Fig. S6A**). The production of PEP by PEPCK is the first and pathway-specific step of gluconeogenesis. Similar to ICL-1, the two isoforms of PEPCK, PCK-1 and PCK-2, were 2-fold elevated in *daf-2* dauers compared to L3 larvae (**Fig. 4B** and **Supplementary Fig. S6A**). Thus, dauers have higher ability to use acetyl-CoA for gluconeogenesis. This ability is further supported by ∼2-fold increase of oxaloacetate producing malate dehydrogenase (MDH-1) and enzymes shared between gluconeogenesis and glycolysis such as enolase (ENOL-1) and aldolase (ALDO-1) (**Fig. 4A, Supplementary Fig. S5** and **S6A**).

As shown above, a simultaneous inactivation of DAF-12 and AAK-2 in *daf-2* at 25°C prevents the gluconeogenic mode and the developmental arrest. Thus, we asked whether DAF-12 or AAK-2, or both, are responsible for the altered expression of the enzymes. To test this, we quantified all 43 enzymes in *daf-2* on DA and *daf-2;aak-2* with or without DA at 25°C. We first analyzed the TCA/glyoxylate cycle branching point. As expected, larvae with high gluconeogenic mode (*daf-2* on DA and *daf-2;aak-2* without DA) showed similar upregulation of ICL-1 as in *daf-2* dauers (**Fig. 4B** and **Supplementary Fig. S5**). Interestingly, *daf-2;aak-2* on DA also had higher amounts of this enzyme despite the low gluconeogenic mode (**Fig. 4B** and **Supplementary Fig. S5**). We reasoned that not only the absolute levels of ICL-1, but its molar ratio in respect to competing TCA cycle enzymes (isocitrate dehydrogenases) controls the isocitrate flow into the glyoxylate shunt (**Fig. 5A**). Indeed, as seen in **Fig. 5B** and **5C**, ICL-1 displayed the lowest molar ratio to all isocitrate dehydrogenase subunits and isoforms in *daf-2* L3 larvae at 15°C. This was consistent with a more intensive TCA cycle. The ratios were overall higher in *daf-2* and *daf-2;aak-2* at 25°C with or without DA (**Fig. 5B** and **5C**). However, in stages with pronounced gluconeogenic mode (*daf-2* without or with DA at 25°C, *daf-2;aak-2* at 25°C) this elevation was much more pronounced (3.5-, 3.8-, and 4.2-fold, respectively) compared to *daf-2;aak-2* at 25°C with DA (2-fold, **Fig. 5B**). In DA-treated *daf-2;aak-2* ICL-1 was much less dominant in respect to the IDHG-1 subunit of the NAD^+^-dependent isocitrate dehydrogenase and, importantly, to the NADP^+^-dependent IDH-1 (**Fig. 5C, Supplementary Fig. S6E and S6F**). Thus, simultaneous inactivation of DAF-12 and AAK-2 lowers the capacity of the glyoxylate shunt.

**Figure 5.**
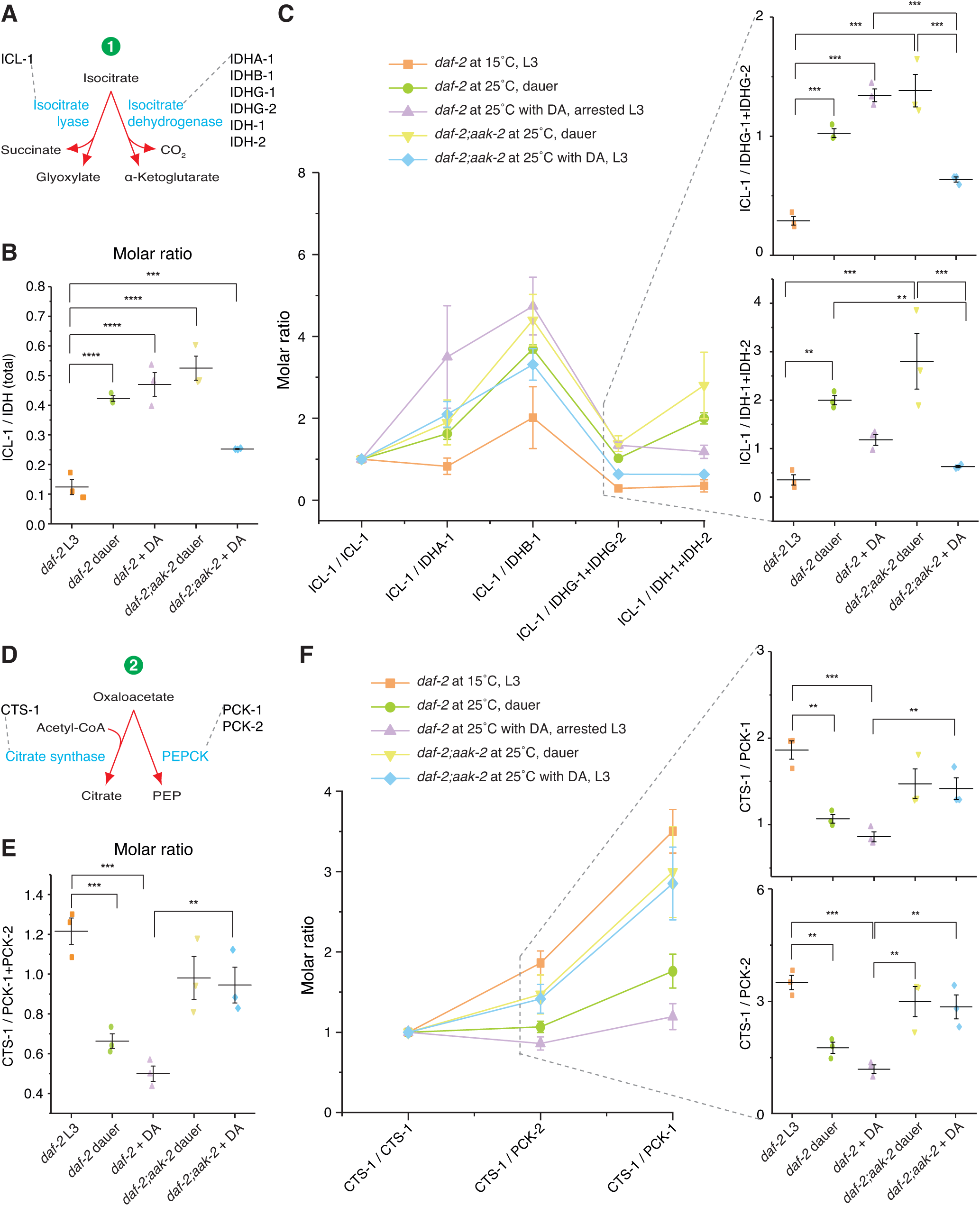
DAF-12 and AAK-2 control the molar ratios of the enzymes at the branching points between competing pathways. (A) Scheme of the first branching point – the entry of isocitrate into glyoxylate or TCA cycle. (B) Molar ratio between ICL-1 and the summed abundances of all isocitrate dehydrogenase isforms and subunits, IDHA-1, IDHB-1, IDHG-1, IDHG-2, IDH-1, and IDH-2, dubbed IDH (total). (C) Molar ratios between ICL-1 and individual isocitrate dehydrogenases. The sums of IDHG-1+IDHG-2 and IDH-1+IDH-2 are provided as a clearer representation due to the low molar abundance of IDHG-2 and IDH-2 compared to IDHG-1 and IDH-1, respectively. Lines between data points are provided for better visualization. (D) Scheme of the second branching point – the recycling of oxaloacetate into citrate or its entry into gluconeogenesis. (E) Molar ratio between CTS-1 and the summed abundances of the two PEPCK isforms PCK-1 and PCK-2. (F) Molar ratio between CTS-1 and the individual PEPCK isforms. Lines between data points are provided for better visualization. In all panels, means ± (S.D.) of 3 biological replicates with 2 technical replicates each; p-values represent p > 0.05 (ns), p ≤ 0.05 (*), p ≤ 0.01 (**), p ≤ 0.001 (***), p ≤ 0.0001 (****). One way ANNOVA was performed with Holm-Bonferroni statistical method.

We next asked whether DAF-12 or AAK-2 control the production of PEP by PEPCK at the second branching point. DA-treated *daf-2* at 25°C had similarly elevated PCK-1 and -2 as *daf-2* dauers (**Fig. 4B and Supplementary Fig. S6A**). The AAK-2 deficient strain, however, showed low PCK-1 and -2 levels regardless of the presence or absence of DA (**Fig. 4B and Supplementary Fig. S6A**). This indicated that AAK-2 might promote conversion of oxaloacetate to PEP (**Fig. 5D**). Accordingly, *daf-2* at 25°C with or without DA showed lower molar ratios of citrate synthase to PEPCK, whereas in *daf-2;aak-2* at 25°C with or without DA they were much higher (**Fig. 5E** and **5F**). Together, our results suggest that both DAF-12 and AAK-2 could induce the switch from TCA to glyoxylate cycle, while AAK-2 promotes the entry of carbon from these cycles into gluconeogenesis.

### The transition from growth to quiescence requires unique constellation of metabolic enzymes

Metabolic switch alters molar ratios of metabolic enzymes in specific way and is not accompanied by a global perturbation of metabolic network. We reasoned that enzymes stoichiometry should be under tighter control than the molar abundances of individual enzymes and, therefore provide a unequivocal phenotype- and context-independent readout of the metabolic state. Indeed, the median coefficient of variation between biological replicates was almost 3.5 fold lower when we calculated it based on the molar ratios between individual enzymes and hexokinase (HXK-3, the first enzyme of glucose utilization) compared to the value obtained by normalization of enzyme abundances to the total protein content (**Fig. 6A**). HXK-3 was chosen because of its low variability (**Fig. Supplementary Fig. S6A**). Hence, the stoichiometry within the enzyme network as a whole (with an emphasis on branching points) may determine the state of metabolism and, thus, development. To test this hypothesis, we subjected the dataset comprised of molar ratios between every enzyme and HXK-3 in *daf-2* at 15°C and *daf-2* and *daf-2;aak-2* with or without DA at 25°C to principle component analysis (PCA). As seen in **Fig. 6B, C,** and **D**, the principal components 1 and 2 of the normalized data (each protein had a mean of 0 and a standard deviation of 1) could easily classify the larval stages according to their developmental state. More specifically, stages of active growth (*daf-2* at 15°C and *daf-2;aak-2* with DA at 25°C) were well separated from arrested stages. To support the validity of the obtained information, we trained a linear discriminate analysis (LDA) classifier (link to the code of the algorithm is available in **Methods**) on the data obtained with L3 (*daf-2* at 15°C) and dauer larvae (*daf-2* at 25°C) and performed ten-fold cross-validation. Astoundingly, using the LDA classifier, we predicted the developmental outcome (growth versus quiescence) of the rest of the samples with a prediction accuracy of 94% despite the low number of quantified enzymes (**Fig. 6E**). Thus, the molar stoichiometry metabolic enzymes is unequivocally associated with the transition from growth to quiescence.

**Figure 6.**
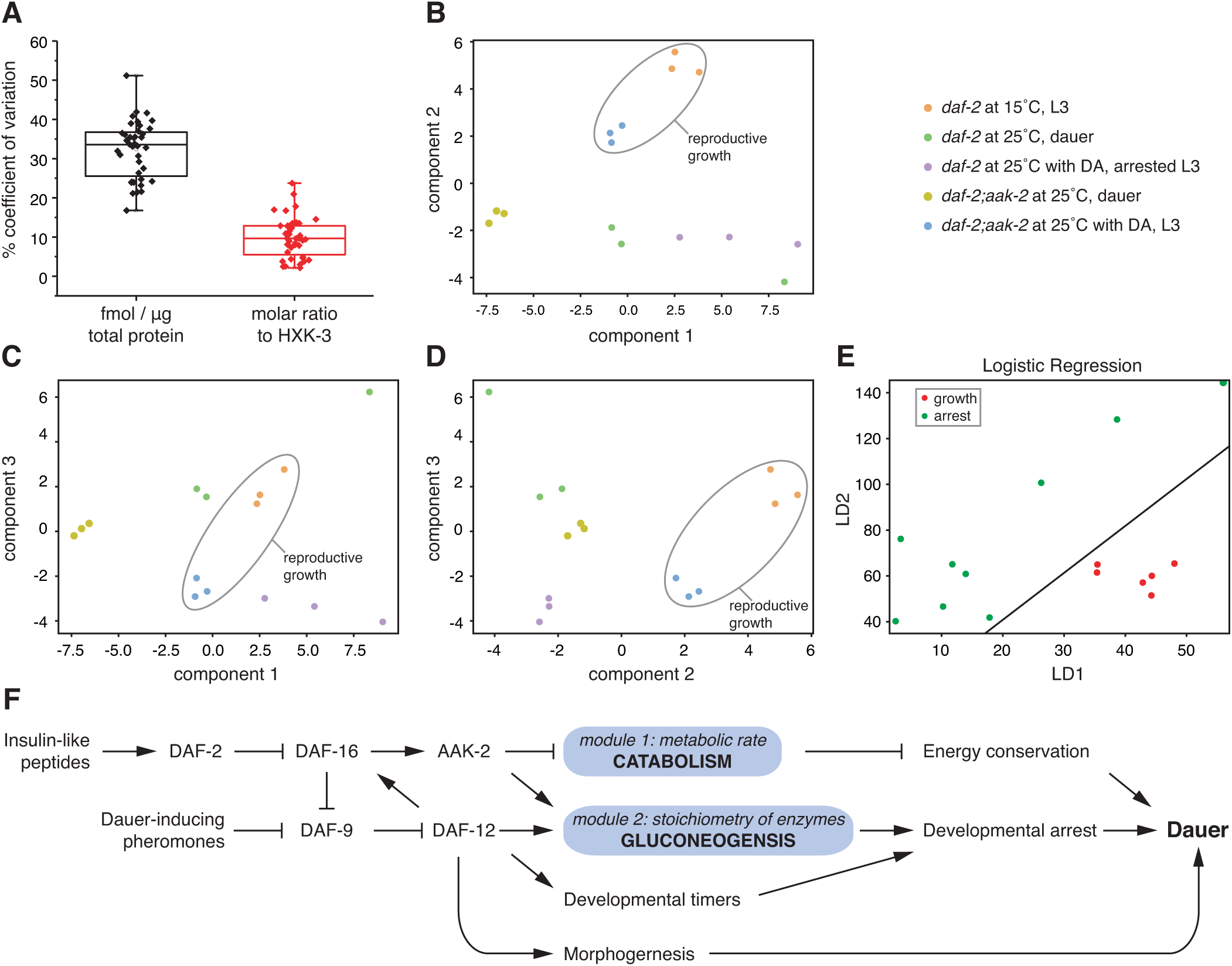
Stoichiometry of 43 enzymes of the central carbon metabolism encodes information for the developmental state. **(A)** Median coefficient of variation between biological replicates of molar abundances of 43 enzymes normalized per total protein content of the sample or expressed as molar ratios to hexokinase (HXK-3). Each point represents the ratio of one protein to HXK-3. The example represents data from L3 larvae (*daf-2* at 15°C). **(B-D)** Principle component analysis (PCA) of molar ratios of 43 individual proteins to hexokinase (HXK-3). Plots represent segregation based on components 1 and 2 (**B**), 1 and 3 (**C**), and 2 and 3 (**D**). Component 1 explains 47.5%, component 2 – 16.1%, and component 3 – 13.6% variance between samples. (E) Lower dimensionality projection of the Linear Discriminant Analysis classifier with a constraint of two components to visualize the results in a 2D-plane. Red dots represent groups in reproductive growth (*daf-2* at 15°C and *daf-2;aak-2* at 25°C with DA). Green dots correspond to arrested larvae of *daf-2* at at 25°C with or without DA and *daf-2;aak-2* at 25°C without DA. (F) Proposed model of metabolic control of the transition to dauer state. Represents the genetic control of metabolic and developmental determinants of dauer formation and their interactions in respect to the establishment of dauer state. The metabolic shift consists of two modules – module 1 comprising the overall metabolic rate, mainly reflecting the catabolism of energy reserves, and module 2 affecting the stoichiometry of metabolic enzymes and, thus, the directionality of metabolic pathways. DAF-16 and AAK-2 inhibit catabolism and promote energy conservation required for the long-term survival of dauers. In parallel to DAF-12, they also promote a shift in the stoichiometry of metabolic enzymes that underlies the enhanced gluconeogenesis and stimulates developmental arrest. The latter occurs precisely at the third larval stage due to the activity of developmental timers, controlled at least partly by DAF-12. The metabolic and physiologic adaptations for survival are complemented by specific morphogenetic program under the control of DAF-12. In (**A**-**E**) - 3 biological replicates with 2 technical replicates each.

## Discussion

This study demonstrates that the transition between growth and quiescence, as well as the mode of metabolism in long-lived insulin signaling mutants, depends on a metabolic switch. The switch is regulated by the concerted action of the insulin, steroid hormone receptor, and AMPK pathways that control key biochemical reactions in the TCA cycle, glyoxylate shunt, and gluconeogenesis. We also show that metabolic mode and morphology are independently regulated in dauer state (**Fig. 6F**), since the arrested larvae of *daf-2;daf-12* and DA-treated *daf-2* mutants display dauer metabolism but reproductive morphology.

Our results provide insight into how major signaling pathways govern the metabolic transition, consisting of two separately controlled modules (**Fig. 6F**). In dauer formation, DAF-16 and AAK-2 inhibit catabolism (module 1) and promote energy conservation, required for the long-term survival of dauers. In parallel, DAF-16, AAK-2, and DAF-12 stimulate gluconeogenesis (module 2) of compounds produced by the glyoxylate pathway. This favors the synthesis of substrates for glycolysis and lowers the TCA cycle and the production of building blocks for anabolism. Consequently, worms cease growth and enter a long-lasting dauer state. When executed in adults, this metabolic mode may contribute to an extension of the lifespan by preventing pathologies associated with activation of growth/reproductive programs during the post-reproductive period [49].

Knowing the molar amounts of enzymes involved in the metabolic switch (in contrast to fold changes between two or more biological conditions) was instrumental in differentiating the contributions of signaling pathways in the metabolic control. By building the first quantitative map of key metabolic enzymes in *C. elegans*, we discovered that, despite the low metabolic rate, dauer larvae maintain an intact metabolic network. Moreover, the enzyme abundance in respect to total protein mass in dauers is even higher than in reproductive larvae. Since catabolic activity and the switch to gluconeogenesis are co-regulated, these findings showcase that the overall metabolic reduction is not achieved through a decrease in the capacity of the metabolic network, but at least partly through suppression of catabolism. Metabolic switch is achieved through adjustments of the enzyme ratios rather than by some on-off mechanism, in which (at least some) enzymes are entirely absent or overproduced. Once put into a favorable environment, dauers could immediately use the available constellation of metabolic enzymes to support reproductive development with no need to re-synthetize them at the cost of massive expenditure of energy.

The exact molar ratios between enzymes acting as metabolic turnouts demonstrated that AAK-2 and DAF-12 equally promoted switching from TCA to glyoxylate cycle, whereas AAK-2 was the primary factor controlling the exit of substrates from the glyoxylate pathway towards gluconeogenesis. Importantly, the combined regulation of the reactions that constitute the two branching points makes the metabolic switch more robust. This is demonstrated by the fact that *daf-2;aak-2* without DA are in gluconeogenic mode despite the high citrate synthase to PEPCK molar ratio. Conceivably, the higher capacity of the glyoxylate shunt in these animals promoted by DAF-12 prevents the carbon from escaping through decarboxylation in the TCA pathway. This leads to higher recycling of oxaloacetate that, although with a lower rate, could enter into gluconeogenesis. However, when DAF-12 is inhibited in *daf-2;aak-2* by addition of DA, the oxaloacetate is not recycled to the same extent and ultimately becomes a substrate of the TCA cycle. Thus, in animals with simultaneously disrupted AAK-2 and DAF-12 signaling, the gluconeogenic mode is not occurring. In addition, the fact that the AMPK and steroid hormone pathways independently govern the switch and have different effects on it shows that the metabolic control is fine-tuned through integration of signaling pathways that work in parallel.

The discovery that the stoichiometry of the enzyme network in the central carbon metabolism predicts the transition from growth to quiescence raises an important question how this information is deciphered. Most probably, signalling cascades are sensitive to steady state concentrations of key metabolites or to fluxes through given pathways. In line with this, we have previously shown that the dauer developmental decision is fine-tuned through regulation of the levels of NADPH required for DA synthesis [14]. This NADPH is generated in reactions directly related to the gluconeogenic mode: the oxidative steps of the pentose phosphate pathway (PPP) and the oxidative decarboxylation of isocitrate by the above-mentioned NADP^+^-dependent isocitrate dehydrogenase IDH-1 [14]. The establishment of the gluconeogenic leads to a reduction of the substrates for NADPH production: the synthesis of trehalose consumes glucose 6-phosphate required for PPP [14], while the glyoxylate pathway competes for isocitrate needed for the IDH-1 reaction. Consequently, when worms switch to gluconeogenic mode, the NADPH levels and thus, the production of DA, are diminished. In the context of the present study, this suggests that the state of gluconeogenesis induces a feedback regulation on the dauer signaling via the steroid hormone pathway. Thus, the second metabolic module directly influences the developmental arrest (**Fig. 6F**). We can predict that future studies will discover further metabolites that influence the development. It must, however, not be excluded that the enzyme concentrations could be also directly sensed, although, to our best knowledge, such system has not been identified yet.

Another challenge will be to understand how the metabolic switch is coordinated with the developmental timing. During growth arrest, cells must simultaneously undergo metabolic depression and acquire dauer-specific fates. Therefore, the master regulators of dauer formation, DAF-12 and DAF-16, should also regulate cell fate decisions. Indeed, DAF-12 has been shown to suppress the progression through larval stages and to determine the cell fates via microRNAs of the *let-7* family serving as developmental timers [15] (**Fig. 6F**), suggesting that it functions as both a metabolic and a developmental “turnout”. However, the fact that the timing of the arrest in the third larval stage is maintained also in the absence of DAF-12 activity suggests that the insulin signaling has similar input via unknown mechanism. In line with this notion, it has been demonstrated that the insulin (via the PTEN homolog DAF-18) and AMPK signaling cascades are crucial for the arrest of the germline cell proliferation during dauer formation [50]. Hence, the investigation of how the synchronization of the metabolic and the developmental programs is achieved will be an important subject for future studies.

Our finding that AMPK modulates the effect of insulin pathway signaling in the control of metabolic mode and growth resolves a long-standing problem in the field. Since class II *daf-2* alleles cannot be rescued by *daf-12* Daf-d mutations or addition of DA [19, 25], it was postulated that an unknown factor mediates larval arrest in worms with active DAF-16 but inactive DAF-12 [25]. Our data indicate that this factor is AAK-2, showing that AMPK signaling has much broader impact on dauer development and metabolism than previously reported. In addition to the known effects of the AMPK on the regulation of energy reserves [41,51,52], we provide evidence that in dauers it also determines the activities of the core metabolic pathways: TCA cycle, gluconeogenesis, glycolysis, and amino acid catabolism. Thus, in a broader context, AMPK in *C. elegans* is not only a modulator of the energy metabolism and mitochondrial function in the responses to metabolic stress [53–58], but also couples the insulin-dependent developmental decisions with the cellular energetic status and nuclear hormone receptor signaling. This notion is consistent with the work of other groups, showing that in the regulation of adult lifespan, AMPK acts at least partially downstream of DAF-16 [59] and *aak-2* loss-of-function alleles suppress the greater longevity of *daf-2* mutants [45]. Indeed, our observation that AAK-2 is required for the full extent of the metabolic switch in long-living adults provides an explanation how the insulin-AMPK interaction promotes longevity at least partly through control of metabolism. Moreover, the FoxO-AMPK-nuclear hormone receptor axis of metabolic regulation might be conserved in higher organisms [60, 61]. Altogether, not only the diapause, but also the lifespan extension are ultimately connected to the combined regulation of metabolic rates and growth, thus highlighting the intricate relationship between growth, development, aging and metabolic state.

## Author contributions

S.P., C.E., J.-M.V., E.K. K.F. and T.V.K. designed the experiments; S.P. conducted phenotypical, fluorescent microscopy and biochemical experiments; B.K.R. and A.S. designed the MS Western analysis which was performed by B.K.R.; S.P., C.E. and J.O. performed microcalorimetry experiments; R.G. optimized and performed CARS microscopy; E.J.M.A. performed bioinformatics analysis; D.V. obtained electron microscopy images; S.P., B.K.R., C.E., J.O., R.G., D.V., A.S., and T.V.K. analyzed the data. All authors discussed the results and S.P. and T.V.K. wrote the manuscript.

## Availability of data and materials

The datasets used and/or analysed during the current study are available from the corresponding author on reasonable request.

## Acknowledgments

We thank all members of Kurzchalia lab for helpful discussions, Prof. Richard Roy for the scientific feedback, and Dr. Iain Patten for the writing support. We are grateful to the Caenorhabditis Genetics Center and to Prof. Adam Antebi for providing worm strains. We thank Prof. Hans-Joachim Knölker for the synthesis of lophanol and dafachronic acid. S.P. is supported by funds from TU Dresden’s Institutional Strategy, financed by the Excellence Initiative of the Federal and State Governments of Saxony and Germany, respectively.

## Methods

### Material and *C. elegans* strains

Lophenol was purchased from Research Plus (Manasquan, NJ USA), [1-^14^C]-acetate (sodium salt) from Hartmann Analytic (Braunschweig, Germany), and Dulbecco’s medium (DMEM) from Invitrogen (Karlsruhe, Germany). (25*S*)-Δ^?^–DA [62–64] and lophanol were produced in the Laboratory of Prof. H.-J. Knölker. All other chemicals were from Sigma-Aldrich (Taufkirchen, Germany).

The Caenorhabditis Genetics Centre (CGC) provided the following *C. elegans* strains: N2 (Bristol strain), *daf-7(e1372), daf-2(e1370), daf-16(mu86), aak-2(gt33), unc-119(ed3);knuSi221.* The strain *unc-119(ed3);knuSi221* contains a single-copy transgenic segment *fib-1*p*::fib-1*(genomic)::eGFP::*fib-1* 3’ UTR + *unc-119*(+). The *E. coli* strain NA22 was also provided by the CGC.

The compound mutant and transgenic strains *daf-2(e1370);daf-12(rh61rh411), daf-16(mu86);daf-2(e1370), daf-2(e1370);aak-2(gt33), daf-2(e1370);knuSi221* (abbreviated *daf-2;fib-1::eGFP*)*, daf-2(e1370);aak-2(gt33);knuSi221* (abbreviated *daf-2;aak-2;fib-1::eGFP*) were generated during this or past studies as published or described below [26, 65].

### Generation of *daf-2;aak-2*

Heterozygous *daf-2(e1370)* males were crossed to *aak-2(gt33)* hermaphrodites. The resulting males were crossed back to the parental *aak-2(gt33)* worms. This gave rise to hermaphrodites that laid eggs at 25°C. A fraction of these eggs developed into dauers. These dauers were shifted to 15°C to re-enter reproductive growth, singled and genotyped by polymerase chain reaction (PCR) for the presence of the *aak-2(gt33)* deletion.

### Generation of daf-2;fib-1::eGFP and daf-2;aak-2;fib-1::eGFP

Males of *unc-119(ed3);knuSi221* were crossed to hermaphrodites of *daf-2(e1370);aak-2(gt33).* Hermaphrodites from the progeny laid eggs at 25°C that developed into dauers. The dauers were left to recover at 15°C and eGFP-positive worms were selected and singled. The progeny of these worms was selected based on fluorescent signal. The presence of *aak-2(gt33)* was tested by PCR and two types of strains were selected as final products of the cross: animals homozygous for the wild-type *aak-2* allele (*daf-2;fib-1::eGFP*) and animals homozygous for *aak-2(gt33)* (*daf-2;aak-2;fib-1::eGFP*).

### Growth and radiolabeling of *C. elegans* strains

The worm strains were routinely propagated on NGM-agar plates complemented with *E. coli* NA22 [66]. When indicated, (25*S*)-Δ^?^–DA was added to the bacteria to a 250 nM final concentration, calculated according to the volume of the NGM-agar. The temperature-sensitive dauer constitutive strains bearing *daf-7(e1372)* or *daf-2(e1370)* alleles were propagated at 15°C, a temperature at which they undergo reproductive growth. To obtain dauers or arrested L3 larvae of these strains, embryos were obtained from gravid adults by hypochlorite treatment [66], incubated overnight at room temperature, and the resulting synchronized L1 larvae were grown at 25°C for 72 hours. The growth on 4-MS-containing medium was performed according to a described method [26]. Briefly, sterol-depleted medium was obtained by substituting agar with chloroform-extracted agarose. NA22 bacteria were grown on a sterol-free DMEM medium, pelleted, rinsed and resuspended in M9 buffer. Two different 4-MS were used depending on the availability: lophenol or lophanol. It must be noted that these two compounds have identical effects to those of dietary sterols [26]. The 4-MS (or cholesterol, when indicated) were added to the bacteria to a 13 *μ*m final concentration according to the volume of the agarose. The worms were propagated on these plates for two consecutive generations. Again, synchronized L1 larvae were grown at 25°C for 72 hours in the second generation until the developmental arrest occurred.

For the microscopy of FIB-1::eGFP in a wild-type background, mixed populations were produced on NGM-agar plates as described above. Reproductive larvae were collected from plates with abundant food and low population density. Dauer larvae were prepared from overcrowded plates, and isolated from other stages by treatment with 1% SDS for 30 minutes followed by separation of the survived dauers from the dead debris of other stages on empty agarose plates on which the dauer larvae quickly dispersed.

To obtain radiolabeled *C. elegans*, the worms were grown on NGM-agar or agarose solid medium (see above), complemented with [1-^14^C]-acetate (sodium salt). The ^14^C-acetate was added to the bacteria and calculated as 0.5 μCi/ml according to the volume of the NGM-agar/agarose.

### Survival, SDS assay and worm sampling for microscopy and biochemical analysis

Dauer(-like) and arrested L3 larvae were prepared by growing the worms on solid medium as described above, collected and washed three times with M9 buffer. The worms were incubated in 15 ml polypropylene centrifuge tubes (Corning, NY, USA) containing 10 ml of autoclaved M9 buffer supplemented with the antibiotics streptomycin (50 μg/ml) and nystatin (10 μg/ml) at 25°C under constant agitation. The density of the population was kept at 500 worms/ml. To monitor the survival, 100 μl aliquots were taken every 2 days and the percentage of live animals was calculated.

For scoring the survival of *daf-2* or *daf-2;aak-2* after SDS treatment, worms were collected from the feeding NGM-agar plates, washed three times with ddH_2_O and resuspended in 10 ml of 1% SDS (w/v) in ddH_2_O in 15 ml polypropylene centrifuge tubes (Corning, NY, USA). After 30 minutes of incubation within the SDS solution at 25°C with shaking, the worms were washed another three times with ddH_2_O and placed on NGM agar plates where the survival was scored. 100 μl aliquots were also used for the preparation of microscopy samples as described below. For biochemical analysis, the worms were washed three times with ddH_2_O, pelleted, snap-frozen in liquid nitrogen and stored at −80°C until further analysis (see below).

### Isothermal microcalorimetry

To measure the heat production during worm development starting from L1 larva onwards, we first purified eggs and plated them on agarose plates without food. After keeping them at 25 °C overnight, synchronized L1 larvae were washed with M9 buffer and diluted to 14.3 worms/μl. 140 μl (∼2000 worms) of each suspension was pipetted into a 4 ml glass ampoule (TA Instruments, New Castle, DE, USA), in which there was already 60 μl of concentrated *E. coli* NA22 in M9 (OD_600_ = 20), so that the starting amount of bacteria was 6 OD_600_. These ampoules were then sealed with aluminium caps equipped with sealing discs (TA Instruments, New Castle, DE, USA).

For the measurements of the heat production of dauer(-like) and arrested L3 larvae, worms were grown on solid medium as described above and washed three times with M9 buffer. 2000 larvae were collected in 200 μl of autoclaved M9 buffer supplemented with the antibiotics streptomycin (50 μg/ml) and nystatin (10 μg/ml) and transferred into 4 ml glass ampoules that were closed with aluminum caps equipped with sealing discs (TA Instruments, New Castle, DE, USA).

Isothermal calorimetric measurements were performed with a TAMIII (Thermal Activity Monitor) instrument (Waters GmbH, Eschborn, Germany) equipped with 12 microcalorimeters in twin configuration (one side for the sample the other for a steel reference) to continuously monitor the metabolic heat produced by *C. elegans* at 25°C for up to 5 days. The samples were held in the TAM III in a waiting position for 15 min before complete insertion followed by 45 min equilibration. In each experiment, thermograms were recorded at least in triplicates. The thermograms represent continuous measurements and no curve fitting was performed.

### Fluorescence and CARS microscopy

For the visualization of FIB-1::eGFP by confocal microscopy and for CARS imaging of lipid deposits, worms were mounted on 2% agarose pads on glass slides (Thermo scientific, Superfrost Plus) and anesthetized with 20 mM sodium azide in M9 buffer. The liquid was aspirated and the pads were covered with cover slips (with 0.17 +/- 0.005 mm cover slips (Menzel-Glaeser). The FIB-1::eGFP was visualized with a Zeiss LSM 880 scanning confocal microscope equipped with a Zeiss i LCI Plan-Neofluar 63x 1.3 Imm Korr DIC objective. eGFP was excited at 488 nm, and fluorescence was detected at the emission band of 490-540 nm. On average, 12 optical sections of 0.09×0.09×1 *μ*m voxel size were collected. To represent the status of the nucleoli in all tissues within the frame, all micrographs are represented as a maximum intensity projection of the Z-stack generated in Fiji.

The imaging of lipid droplets was performed by coherent anti-Stockes Raman scattering (CARS) microscopy [67]. Autogenous two-photon excited fluorescence (TPEF) and second harmonic generation (SHG) optical signals were simultaneously acquired. TPEF was used to differentiate between lipid droplets and autofluorescent lysosome-related organelles [68]. SHG displays collagen type I and was used as reference for anatomical details, e.g. the position of the pharynx [69]. CARS, TPEF and SHG were detected using a multiphoton scanning microscope coupled with two near-infrared picosecond fiber lasers. The optical microscope was an upright Axio Examiner Z.1 equipped with a laser scanning module LSM 7 (all from Carl Zeiss Microscopy GmbH, Jena, Germany) and multiple detectors in non-descanned configuration. The excitation for TPEF and SHG was provided by an Erbium fiber laser (Femto Fiber pro NIR from Toptica Photonics AG, Gräfelfing, Germany) emitting at 781 nm with pulse length of 1.2 ps and maximum emitted power at the source of 100 mW. The TPEF signal in the spectral range 500-550 nm was acquired in reflection. The SHG signal was acquired in transmission mode with band pass (BP) filter (390 ± 9) nm. A second laser source was used to excite the CARS signal. This source (Femto Fiber pro TNIR from Toptica Photonics AG) is tunable in the range 850 - 1100 nm and has a pulse length of 0.8 ps. In all CARS experiments the wavelength was set to 1005 nm (emitted power at the source:1.5 mW), to resonantly excite the symmetric stretching vibration of methylene groups at 2850 cm^-1^. The CARS signal was collected in transmission mode and selected using a BP filter (640 ± 7) nm. A water immersion objective W Plan-Apochromat 20×/1.0 (Carl Zeiss Microscopy GmbH) was used. Due to the transmission of optical elements, the laser power in the sample was 52 mW. CARS, TPEF and SHG were combined as RGB images (red: CARS; green: TPEF; blue: SHG). An automatic tiling procedure enabled by the microscope software ZEN was used for acquisition of images larger than the field of view of the microscope objective.

### High-pressure freezing and electron microscopy

Worms were directly frozen without any additives with a high-pressure freezing unit (EMPACT2, Leica), followed by automated freeze substitution (AFS2, Leica) in acetone cocktail (containing 1% osmium tetroxide, 0.1% uranyl acetate and 0.5% glutaraldehyde), with a slope of 3.0°C/hour, from −90°C up to 0°C (including a rest for 15 hours at −30°C). At room temperature, samples were rinsed with acetone and stepwise infiltrated with mixtures of acetone and LX112-resin (Ladd Research) from 1/3 over ½ to 2/3 the amount of resin (1.5 hours each step). Samples were left in pure resin overnight, then for another four hours in fresh resin before mounting them between slides and polymerizing at 60°C. Transverse sections (70 nm) were taken with an ultramicrotome (Ultracut UCT, Leica), and post-contrasted in 1% uranyl acetate in 70% methanol followed by lead citrate. The sections were examined under electron microscope (Philips Tecnai12, FEI) at 120 kV, and photographs were taken with a TVIPS-camera (Tietz).

### Organic extraction and thin layer chromatography

Frozen worm pellets were homogenized by three rounds of thawing in an ultrasonication bath and freezing, and extracted using a standard method [70]. In all experiments, the samples consisted of similar numbers of worms (∼20 000 larvae). After phase separation, lipids and hydrophilic metabolites were recovered from the organic and aqueous phases, respectively. Non-radioactive samples were normalized for the number of worms. Radioactive ^14^C-acetate-labeled samples were normalized for the number of worms to visualize the rate of catabolism of TGs, phospholipids, trehalose and amino acids in *daf-2;aak-2* or according to the total radioactivity to determine the state of the gluconeogenesis in different worms strains. In the latter case, the normalization method was chosen to obtain information on the relative abundance of the various metabolites.

TLC was performed on 10 cm HPTLC plates (Merck, Darmstadt, Germany). The running system for sugar detection was chloroform-methanol-water (4:4:1, v/v/v) and chloroform-methanol-water (45:18:3, v/v/v) for phospholipid detection. 2D-TLC for the visualization of hydrophilic metabolites was done using 1-propanol–methanol–ammonia (32%)–water (28:8:7:7, v/v/v/v) as 1^st^ system and 1-butanol–acetone–glacial acetic acid–water (35:35:7:23, v/v/v/v) as the 2^nd^. The TLC plates were sprayed with Molisch reagent for sugar detection, with ninhydrin for visualization of amino acid, and with 3% copper (II) acetate in 10% orthophosphoric acid for imaging of TGs and phospholipids. TLC plates containing radioactive samples were sprayed with EN^3^HANCE spray surface autoradiography enhancer (Perkin Elmer, Waltham, MA, USA) and exposed to X-ray film (Kodak Biomax MR, Sigma-Aldrich, Taufkirchen, Germany). The X-ray films were scanned and the band intensities of TGs, phosphatidylethanolamines and trehalose were calculated in Fiji by determining the corresponding optical density peak areas.

### MS Western absolute quantification of metabolic enzymes

Absolute protein quantification was performed using MS Western [48]. All worm strains were washed twice with M9 buffer, counted, collected and snap frozen in liquid nitrogen for later analysis. The frozen worms were thawed on ice and crushed using a micro hand mixer (Carl Roth, Germany). The crude extract was then centrifuged for 15 min at 13000 rpm, 4°C to remove any tissue debris. The clear supernatant was then transferred to a fresh Protein Lo-Bind tube (Eppendorf, Hamburg, Germany). Total protein content of the samples were estimates using BCA assay (Thermo scientific, Germany) and 60 μg (∼3500 worms) of total protein content was loaded on to a precast 4 to 20% gradient 1-mm thick polyacrylamide mini-gels were from Anamed Elektrophorese (Rodau, Germany) for 1D SDS PAGE. Separate gels were run for 1 pmol of BSA and isotopically labelled lysine (K) and arginine (R) incorporated chimeric standard containing 3-5 unique top N quantitypic peptides from 53 metabolic enzymes spanning glycolysis, gluconeogenesis, TCA cycle and glyoxylate shunt and 5 peptides from BSA for quantifying the standard [48]. Undetectable proteins or proteins without detectable unique sequences like GPD-1, GPD-3, HXK-1, ALH-4, ALH-5, ALH-11, ALH-2, SODH-2, SUCL-1, and SDHD-1 were not included in this analysis. Peptides containing methionine and cysteine were excluded as the former can be variably oxidised and the later can form disulphide bridges. The sample was cut into 6 gel fractions and each fraction was co digested with BSA and the chimeric standard using Trypsin Gold, mass spectrometry grade, (Promega, Madison). Mass spectra was acquired in data-dependent acquisition mode in a Q-Exactive HF (Thermo Scientific, Bremen, Germany) coupled with a Dionex Ultimate 3000-HPLC system (Thermo Scientific, Bremen, Germany). Peptide matching was carried out using Mascot v.2.2.04 software (Matrix Science, London, UK) against *Caenorhabditis elegans* (November 2016) proteome downloaded from Uniprot. A precursor mass tolerance of 5ppm and fragment mass tolerance of 0.03 Da was applied, fixed modification: carbamidomethyl (C); variable modifications: acetyl (protein N terminus), oxidation (M); labels: ^13^C(6) (K) and ^13^C(6)^15^N(4) (R); cleavage specificity: trypsin, with up to 2 missed cleavages allowed. Peptides having the ions score above 15 were accepted (significance threshold p < 0.05). The chromatographic alignment and feature detection were carried using Progenesis LC-MS v.4.1 (Nonlinear Dynamics, UK). The absolute quantification was performed by calculating the abundances for the labelled and the unlabelled peptide using an in-house software.

### Principal component analysis and linear discriminant analysis

Molar ratio of all 43 enzymes to HXK-3 from all 5 conditions were used as the data space for further analysis. The initial data exploration was carried out with PCA. Most of the data variance was explained with first 3 components. Plotting the data, it was linearly dividable in the lower dimension 2D projection.

We then chose to make a classifier using Linear Discriminant Analysis (LDA) owing to the small size of the data set. The data was scaled to each feature and it fitted a normal distribution. To validate the performance of the classifier, we used 10-fold cross validation. The data was split into 10 parts and one part was used to test the data trained on rest 9 parts. An accuracy of 94% was achieved. To visualize the classified data, we used a logistic regression to define areas on a projected 2D-surface and the points on the surface represent the samples used in the classification.

The code used for classification and visualization of the data can be found in the link (< placeholder https://cloud.mpi-cbg.de/index.php/s/15V5MqAZlptnX8I >) The Jupyter notebook loads the data as a pandas data frame and performs principal component analysis as well as Linear Discriminant Analysis, for data exploration and classification. For visualization a logistic regression was used to project the higher dimensionality data into a lower 2D-plane.

---

**Supplementary Figure S1.**
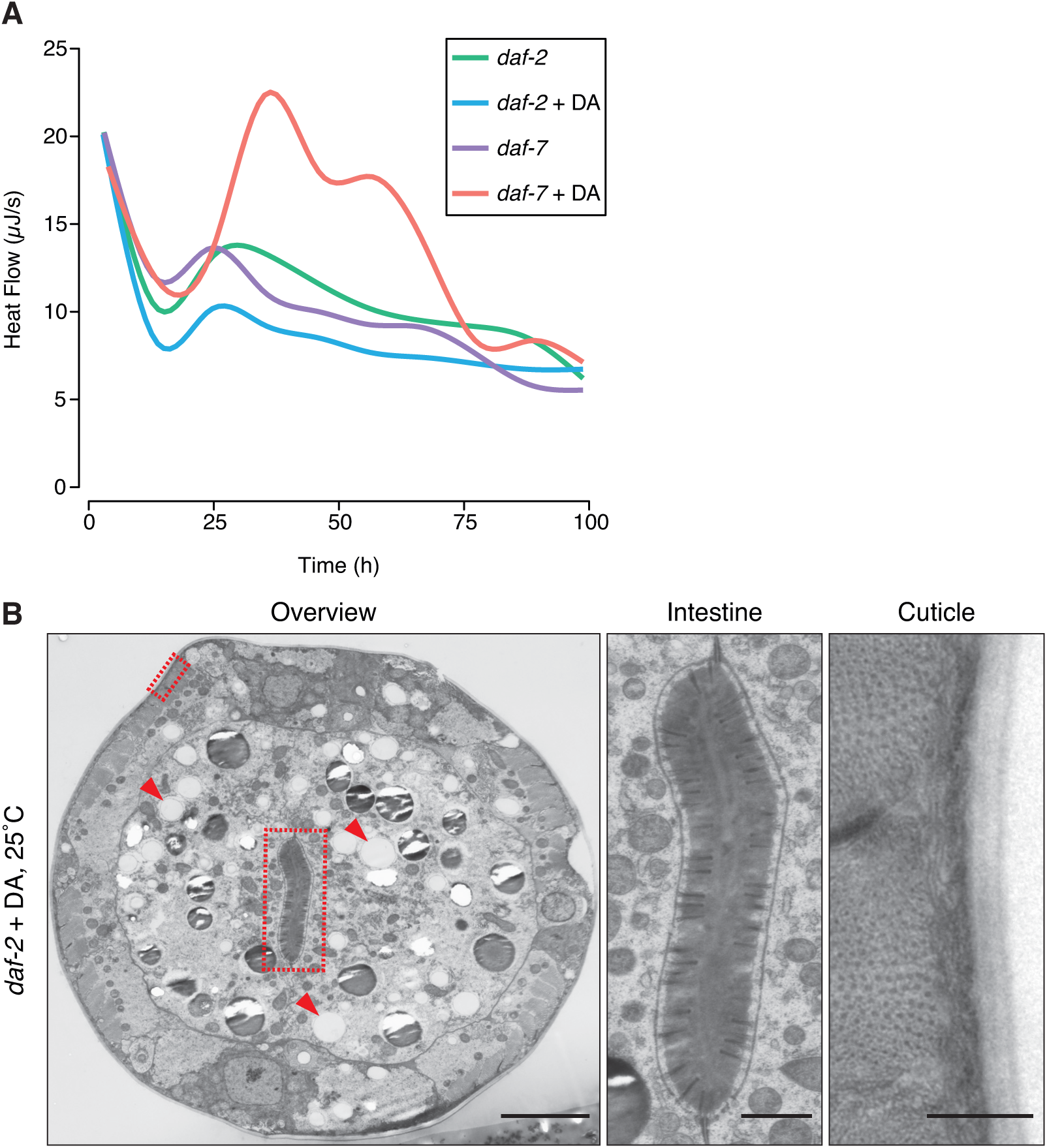
Effect of DA on the heat production of *daf-2* and *daf-7* and the morphology of *daf-2*. (**A**) Heat flow of *daf-2* and *daf-7* grown at 25°C in the presence or absence of DA. Inhibition of DAF-12 suppresses the switch to low heat production in *daf-7* but not in *daf-2*. (**B**) Electron micrograph of a *daf-2* arrested L3 larva grown at 25°C in the presence of DA. The body is not radially constricted but multiple lipid droplets are visible (left panel, arrowheads). Alae are absent (left panel). The gut lumen is elongated with multiple microvilli (central panel, big rectangle on left panel) and the cuticle has no striated layer (right panel, small rectangle on left panel). In (**A**), n=2 for each condition. DA – dafachronic acid. In (**B**), representative images of five worms. Scale bars 5 *μ*m (left panel), 1 *μ*m (central panel) and 0.5 *μ*m (right panel).

**Supplementary Figure S2.**
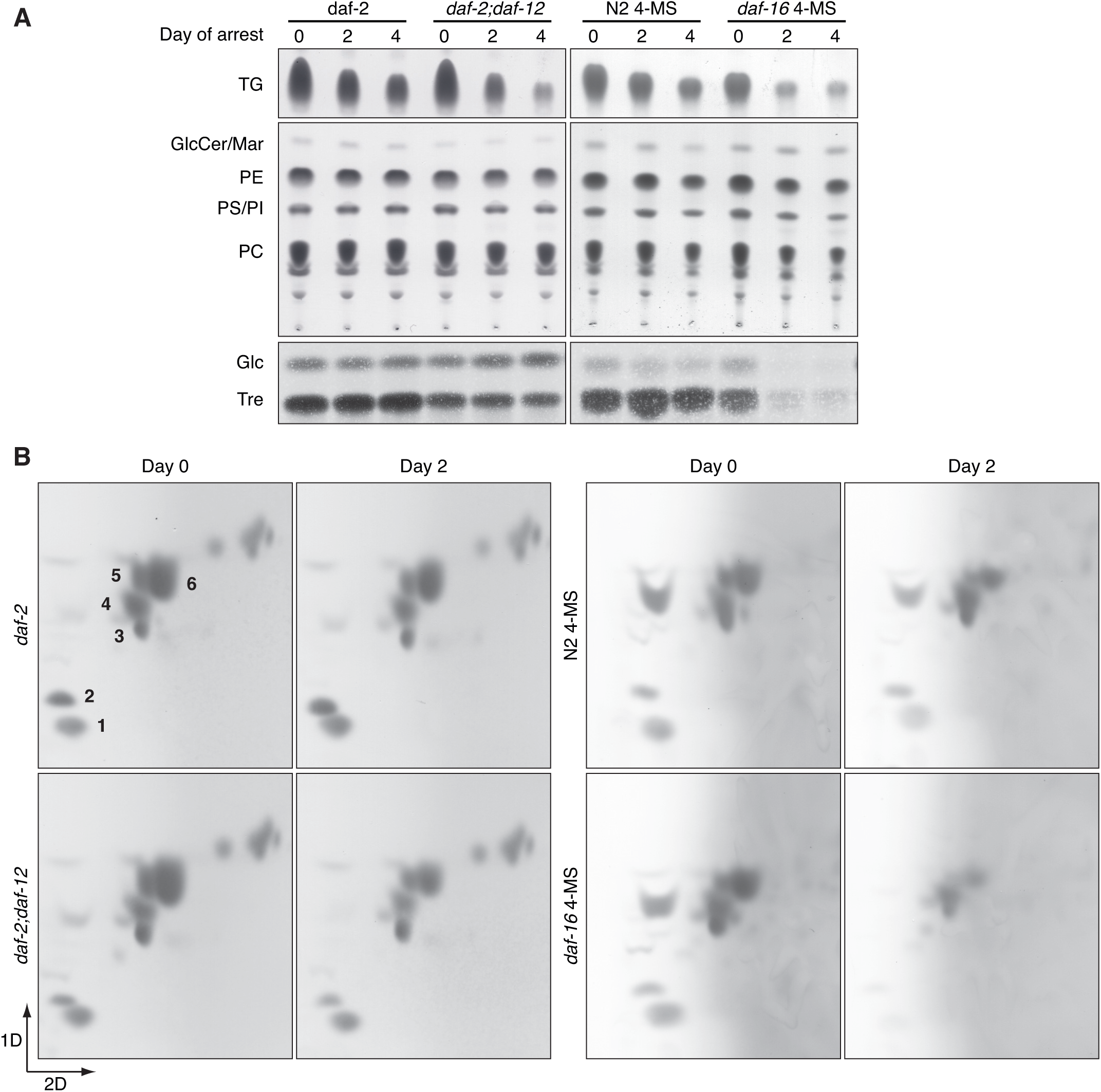
DAF-16 controls the catabolism of energy reserves in dauer larvae. **(A)** TLC of lipids and sugars in *daf-2* dauers and *daf-2;daf-12* arrested L3 larvae grown at 25°C, and wild-type (N2) dauers and *daf-16* dauer-like larvae grown on 4-MS in the period after the developmental arrest is completed. While the *daf-2*, *daf-2;daf-12* and wild-type animals show conservation of TGs, trehalose and glucose, in *daf-16* only traces of these compounds are visible within 2 days of arrest. This is not the case with phospholipids, which are preserved to a good extent in all larvae. **(B)** 2D-TLC of amino acids from the same types of animals as in (**a**). *daf-2*, *daf-2;daf-12* and wild-type animals are able to preserve the bulk amino acids, while in *daf-16* larvae after 2 days of arrest the amino acid levels are very low. TG – triglycerides, GlcCer – glucosylceramides, Mar – maradolipids, PE – phosphatidylethanolamine, PS – phosphatidylserine, PI – phosphatidylinositol, PC – phosphatidylcholine, Glc – glucose, Tre – trehalose, 1 – arginine, 2 – lysine, 3 – glutamate, 4 – glycine/serine, 5 – glutamine, 6 – alanine/threonine. Representative images of at least 2 experiments.

**Supplementary Figure S3.**
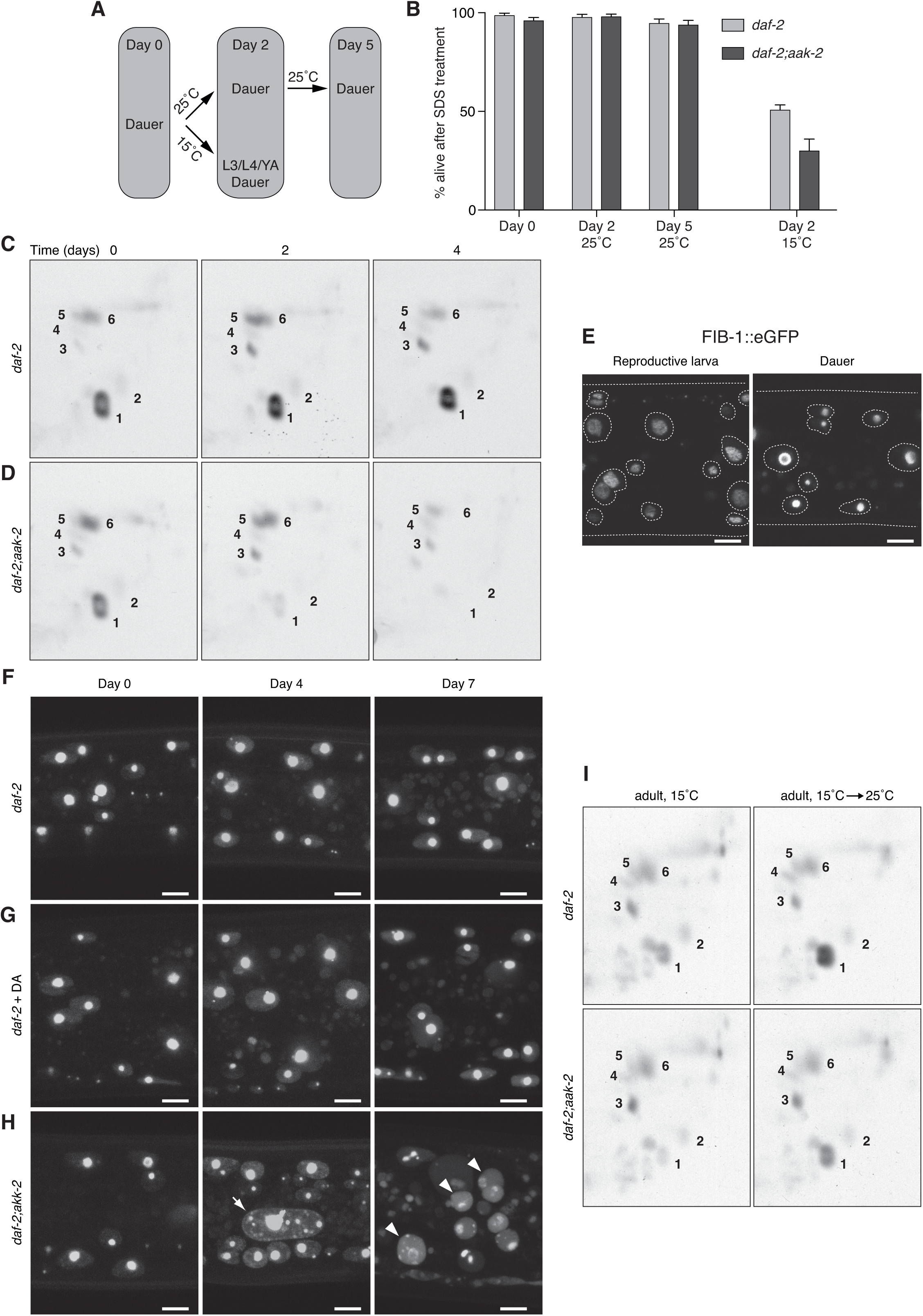
AAK-2 does not inhibit exit from dauer state but is required for preservation of energy reserves in dauers and for gluconeogenic mode in adults. **(A)** Scheme of the SDS treatment experiment. Dauer larvae of *daf-2* and *daf-2;aak-2* were obtained by incubation at the restrictive temperature (25 °C) and subjected to treatment with 1% SDS immediately after the completion of the dauer formation (Day 0) or after prolonged incubation at 25 °C (Day 2 and 5) in the presence of food. After SDS treatment, the survival of the animals was scored. Separately, dauer larvae were allowed to exit by a shift to permissive temperature (15 °C) until day 2, when a mixture of reproductive stages (L3, L4 larvae and young adults (YA)) and not recovered dauers was formed. This population was also treated with SDS. **(B)** Survival after SDS treatment. Both *daf-2* and *daf-2;aak-2* larvae show almost 100% survival even after five days at the restrictive temperature (25 °C). In contrast, the worms shifted to permissive temperature (15 °C) display substantial sensitivity to SDS. **(C)** 2D-TLC of ^14^C-acetate labelled sugars and amino acids from *daf-2* dauers measured at different time points after the arrest. The depicted compounds are well preserved over time. **(D)** 2D-TLC of ^14^C-acetate labelled sugars and amino acids from *daf-2;aak-2* dauers. Unlike *daf-2*, *daf-2;aak-2* are depleted of sugars and amino acids very fast. **(E)** FIB-1::eGFP localizes to nucleoli in both reproductive and dauer larvae. The outlines of the nuclei are indicated by dashed lines. **(F)** *daf-2;fib-1::eGFP* dauers at different time points after dauer arrest. FIB-1 is localized to the nuclei with highest concentration in the nucleoli in all cells. Over time, this localization is retained. **(G)** *daf-2;fib-1::eGFP* arrested L3 larvae grown on DA at different time points after the arrest. Over time, the animals maintain nucleolar localization of FIB-1. **(H)** *daf-2;aak-2;fib-1::eGFP* dauers at different time points after arrest. Early after arrest, FIB-1 is localized to the nucleoli. After 4 days, FIB-1 is still detected in the nucleoli but also in multiple smaller granules dispersed in the nucleoplasm of some cells (arrow). After 7 days, FIB-1 is almost completely dissolved in the nucleoplasm of most of the cells (arrowheads). **(I)** 2D-TLC of ^14^C-acetate-labelled metabolites from *daf-2* and *daf-2;aak-2* adults grown at 15°C (left panels) or switched from 15°C to 25°C after L4 stage (right panels). In (**B**), means + SD of 2 experiments performed in triplicates. In (**C**) and (**D**), and (**I**) representative images from 2 experiments. 1 – trehalose, 2 – glucose, 3 – glutamate, 4 – glycine/serine, 5 – glutamine, 6 – alanine/threonine. In (**E-H**), maximum intensity Z-projection of the eGFP fluorescence. DA-dafachronic acid. Scale bars – 5 *μ*m. Representative images of 2 experiments with at least 7 animals (**E**) and 3 experiments with at least 7 animals (**F**, **G**, and **H**).

**Supplementary Figure S4.**
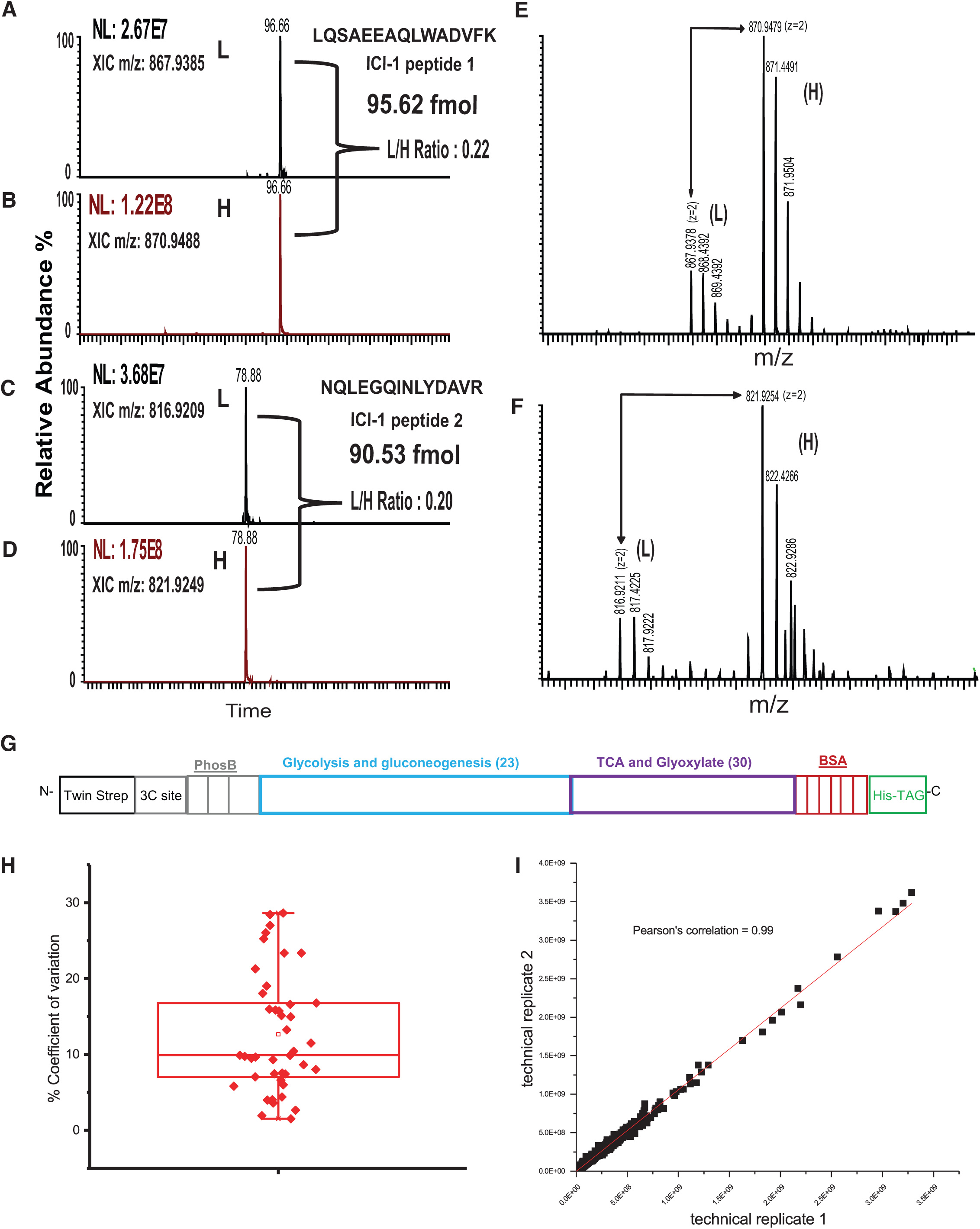
LC-MS/MS (MS Western) analysis of metabolic enzymes. (**A**-**F**) Multiple peptide based concordant quantification of an example Protein Isocitrate lyase (ICL-1). Extracted ion chromatograms (XIC) of ICL-1 endogenous peptides **(A)** LQSAEEAQLWADVFK and **(C)** NQLEGQINLYDAVR their corresponding co-eluting labelled peptides **(B** and **D)** from artificial chimeric standard. **(E)** and **(F)** show the isotopic distribution of the light (L) and heavy (H) peptides. The light to heavy ratio (L/H) are similar for both peptides. The quantification is performed by comparing the peak abundances of known amount of the chimeric standard to the peak abundances of the endogenous peptides to calculate the amount in fmole. The final amount is reported as an average of the calculated values for all peptides. (G) Scheme of the chimeric construct used in the MS Western measurements. (**H**) Coefficient of variation (CV %) distribution of 43 proteins were each point represents one protein. Proteome of L3 larvae (*daf-2* at 15°C is given as example) The CV % was calculated for each protein in one sample by the following formula, 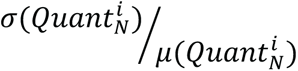 were *i* represents the protein, N represents the number of quantitypic peptide in protein (*i)* and the *Quant* is the amount in fmole. *σ* and *μ* represents the standard deviation and mean respectively. The median CV was less than 10 % (9.336 % ± 5.3%). (**I**) Scatter diagram demonstrating the concordance between technical replicates within the whole proteomics data set of L3 and dauer (*daf-2* at 15°C and 25°C).

**Supplementary Figure S5.**
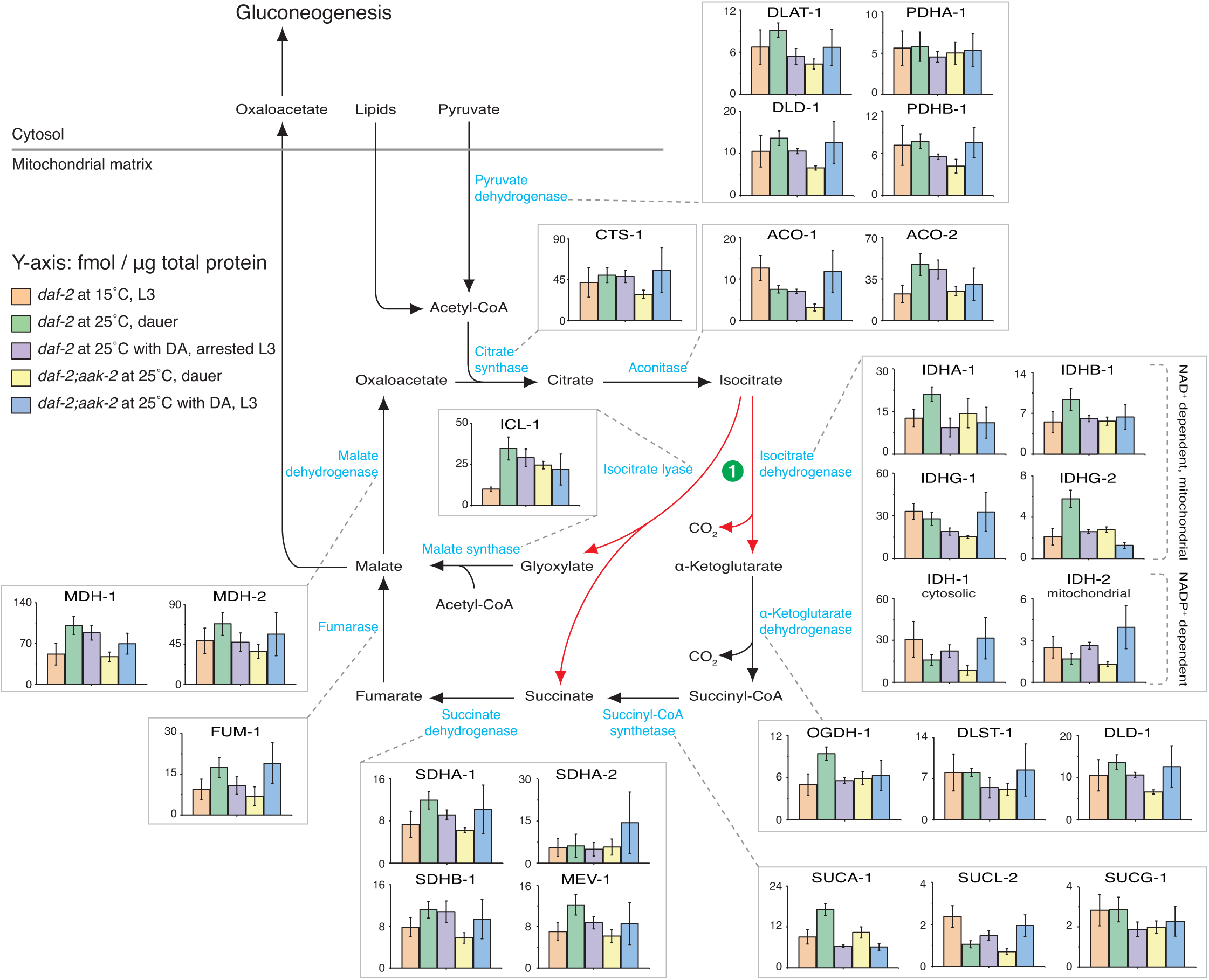
Control of the TCA and glyoxylate cycle. Absolute quantification of enzymes of the TCA cycle and the glyoxylate shunt in *daf-2* and *daf-2;aak-2* grown at 25°C with or without DA compared to *daf-2* animals at 15°C. The red arrows and the green circle represent the two competing reactions at the point of divergence between TCA and glyoxylate pathway. Means ± standard deviation (S.D.) of 3 biological replicates with 2 technical replicates each.

**Supplementary Figure S6.**
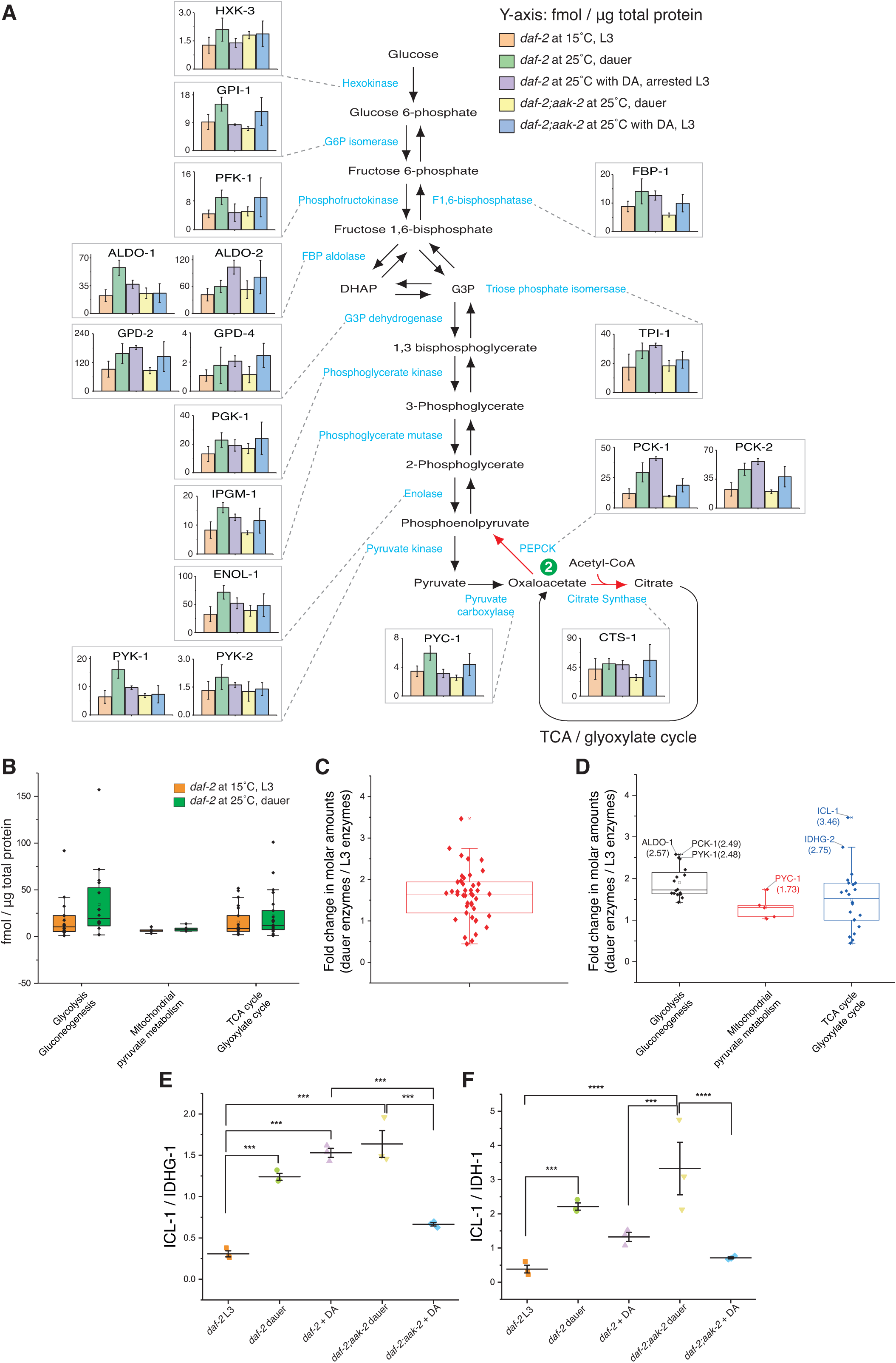
Control of the gluconeogenesis and the molar ratios between ICL-1 and individual isocitrate dehydrogenases. **(A)** Absolute quantification of enzymes of the gluconeogenesis and glycolysis in *daf-2* and *daf-2;aak-2* grown at 25°C with or without DA compared to *daf-2* animals at 15°C. The red arrows and the green circle represent the two competing reactions at the point of divergence between oxaloacetate recycling and entry into gluconeogenesis. **(B)** Median molar abundance ± S.D. of metabolic pathways. **(C)** Median fold change ± S.D. of molar abundances of all proteins in dauers compared to L3 larvae. **(D)** Median fold change ± S.D. of molar abundances of proteins according to the metabolic pathways in dauers compared to L3 larvae. **(E)** Molar ratio between ICL-1 and IDHG-1. **(F)** Molar ratio between ICL-1 and IDH-1. In (**A**), data is represented as means ± standard deviation (S.D.) of 3 biological replicates with 2 technical replicates each. In (**B**), (**C**), and (**D**), median values ± standard deviation (S.D.) of 3 biological replicates with 2 technical replicates each. In (**E** and **F**), means ± (S.D.) of 3 biological replicates with 2 technical replicates each; p-values represent p > 0.05 (ns), p ≤ 0.05 (*), p ≤ 0.01 (**), p ≤ 0.001 (***), p ≤ 0.0001 (****). One way ANNOVA was performed with Holm-Bonferroni statistical method.

